# Tarloxotinib targets tumor hypoxia to improve therapeutic efficacy of immune checkpoint inhibition in a TLR9 dependent manner

**DOI:** 10.1101/2025.02.24.635378

**Authors:** Zhe Fu, Victoria Jackson-Patel, Kathryn J. Farrand, Chris P. Guise, Shevan Silva, Alisha Dabb, Xiaojing Lin, Emily Liu, Amir Ashoorzadeh, Jeff B. Smaill, Ian F. Hermans, Adam V. Patterson

## Abstract

Immune checkpoint inhibitors can elicit deep and durable immune responses although most cancer patients fail to experience long-term remission. There is consensus that tumor hypoxia coordinates a multitude of underlying resistance mechanisms that contribute to treatment failure. Here we show that the clinical-stage hypoxia-activated prodrug tarloxotinib lowers tumor hypoxia 8-fold in the EGFR-dependent MB49 syngeneic tumor model, attracting a effector CD8^+^ T cell infiltrate into an oxygen enriched tumor microenvironment, leading to potentiation of checkpoint inhibitor activity. Various methodologies, including CD8^+^ T cell depletion and exogenous T cell priming confirmed that tarloxotinib has a positive impact on the function of activated CD8^+^ T cells. Whilst anti-PD-L1, anti-PD-1 and anti-CTLA-4 treatments all benefited from the remodeled tumor microenvironment, anti-PD-L1 was most responsive to tarloxotinib coadministration, providing a 100% complete response rate and >360% improvement in tumor growth delay (day 19; p<0.0001). This robust interaction was associated with MB49 tumor enrichment of CD8^+^ T cell infiltrate from 8% to 43% of the total CD45^+^ population. To further understand these promising observations, comparative RNA transcript analysis of 770 cancer/immune related genes in naïve and tarloxotinib treated tumors highlighted the strong induction of gene clusters related to co-stimulatory signaling, cytokine/chemokine signaling, interferon signaling, immune cell adhesion/migration, antigen presentation and the lymphoid/myeloid compartments. Serum protein analysis confirmed upregulation of a mixture of cytokines/chemokines, including IL-6, IL-12p40, IFNγ, G-CSF and MIP-1β, amongst others. The source of this broad immunogenic response was identified as toll-like receptor 9 (TLR9) dependent, with the positive interaction between anti-PD-L1 and tarloxotinib blocked in MyD88 or TLR9 knockout mice. Further, the observed therapeutic interaction was still evident in MC38 and EG7.OVA syngeneic tumor models, both refractory to tarloxotinib by virtue of EGFR-signal independence. Consistent with the clinical experience of minimal systemic toxicities related to EGFR inhibition (e.g. diarrhea), mouse body weight loss was minimal across all *in vivo* studies and histopathology screening for evidence of lung fibrosis proved negative. Tarloxotinib therefore represents a systemically administered small molecule with an established clinical safety profile that is capable of activating TLR9 signaling within tumors whilst remodeling the microenvironment to facilitate efficacy of checkpoint blockade.

**What is already known on this topic:** Solid tumor resistance to immune checkpoint inhibitors (ICI) is orchestrated through a diverse collection of hypoxia-driven mechanisms. In preclinical models modifying the tumor microenvironment (TME) to lessen hypoxia typically improves responses to ICI.

**What this study adds:** Tarloxotinib is a hypoxia-activated prodrug (HAP) of an irreversible EGFR/HER2 TKI that profoundly remodels the TME, both eliminating tumor hypoxia and elevating cytokine/chemokine production via a TLR9 dependent effect. Together, this results in a marked tumor influx of activated CD8^+^ T cells (and other TILs), that results in major improvements in the efficacy of ICIs, particularly anti-PD-L1.

**How this study might affect research, practice or policy:** Tarloxotinib is the first example of a tumor-targeted small molecule TLR9 stimulant that restores ICI sensitivity in a range of syngeneic tumor models. Phase I/II data has demonstrated that tarloxotinib is well tolerated, with few EGFR-dependent (on-target) side-effects, reflecting minimization of normal tissue exposure with the tumor-targeted HAP approach. Importantly, it also optimizes tumor selective exposure, offering a compelling clinical rationale for evaluation of this combination in cancer patients experiencing ICI relapse or resistance.

## INTRODUCTION

Immune checkpoint inhibitors (ICI) have revolutionized oncology practice with significant clinical impact in several cancer indications. Typically administered as blocking antibodies, ICI interrupt interactions between inhibitory checkpoint receptors expressed by activated T cells and their respective ligands expressed on antigen-presenting cells, or cells within the tumor microenvironment (TME). This prevents negative regulation of a weak anti-tumor response that often develop spontaneously in patients (1). To date, inhibitors of cytotoxic T-lymphocyte antigen-4 (CTLA-4), programmed death protein-1 (PD-1) and its ligand programmed death-ligand 1 (PD-L1), and lymphocyte-activation gene 3 (LAG-3) have been approved for clinical use against a variety of cancers, either as a monotherapy or in combination with other treatment modalities. Although capable of unleashing robust and durable anti-tumor immune responses, even against advance-staged cancers, patients that gain long-term benefits still remain in the minority, suggesting underlying mechanisms of resistance (1, 2, 3). Treatment failure is multifaceted, involving tumor intrinsic and extrinsic factors, including a lack of suitable or sufficient tumor antigens for T cell recognition, defects in presentation of tumor antigens, dysfunction of the T cell response due to chronic stimulation or exposure to suppressive factors, physical barriers limiting T cell infiltration, negative influence of the microbiome, and host factors such as genetics, sex and age that affect host immunity (3, 4, 5). To improve outcomes to ICI, it is therefore necessary for the immune response to overcome these immunosuppressive mechanisms to elicit a robust and durable anti-tumor response.

Hypoxia is a common feature of the TME within solid tumors, contributing to many of the suppressive mechanisms that present a critical barrier to effective anti-tumor immunity (3, 6). Regions of tumor hypoxia are associated with recruitment of immunosuppressive cells including regulatory T cells (7, 8), myeloid-derived suppressor cells (MDSC) and tumor-associated macrophages (9, 10), while T cells are often excluded (11). The function of effector T cells and dendritic cells (DC) is compromised through adenosine accumulation (12), while hypoxia-inducible factor (HIF)-1α expression leads to upregulation of inhibitory molecules such as CTLA-4 and PD-L1 (13, 14). Strategies have been employed to reduce tumor hypoxia to improve anti-tumor immune responses, with some showing promising therapeutic benefits including hyperbaric oxygen therapy (15, 16), treatment with metformin to reduce cellular oxygen consumption (17), and use of HIF-1α inhibitors (18). Hypoxia-activated prodrugs (HAP) are a class of drugs that exploit the low oxygen levels within tumors to become active through metabolic conversion, achieving high intratumoral doses of anticancer agents with significantly reduced off-target effects (11, 19, 20). Consequently, regions of hypoxia are preferentially targeted and eliminated, potentially removing the driver of many immunosuppressive pathways. Use of HAP may therefore combine effectively with ICI, leading to improved outcomes. In fact, a recent study with the DNA alkylator HAP evofosfamide demonstrated remodeling of the TME which improved CD8^+^ T cell effector function in murine prostate and lung cancer models (11, 21), and in patients, evofosfamide resensitized immune-refractory prostate, head-and-neck and pancreatic tumors to ICI therapy (22).

Here we investigated whether responses to ICI could be improved by combining treatment with tarloxotinib, a HAP that releases tarloxotinib-E, a potent irreversible tyrosine kinase inhibitor (TKI) of the epidermal growth factor receptor (EGFR)/human epidermal growth factor receptor-type 2 (HER-2) (**Figure 1A**). Tarloxotinib has shown single-agent activity in various EGFR/HER2-dependent xenograft models (23, 24), and clinical studies demonstrated single-agent activity in relapsed/refractory HER2 exon20ins non-small cell lung cancer (20, 25). It has a excellent clinical safety profile due to its stringent hypoxia-selective mechanism that releases payload only within the severe hypoxia regions typically present in the TME. Because tarloxotinib releases a EGFR/HER2 inhibitor, rather than a broad genotoxic DNA-damaging agent, it was anticipated that undesirable effects on immune cells would be avoided, potentially making it an ideal partner for ICI.

**Figure 1.**
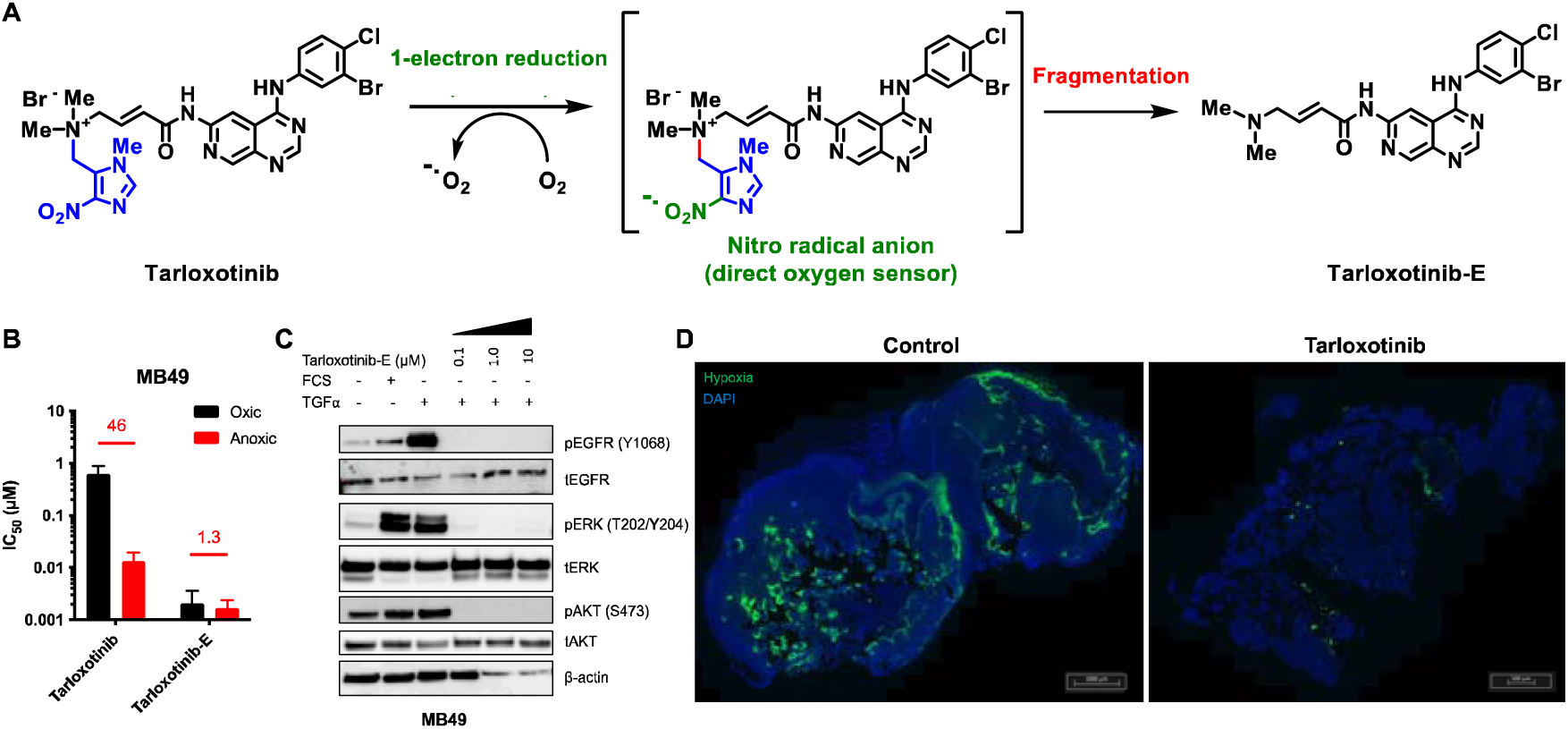
Tarloxotinib-E demonstrates on-target inhibition of EGFR signaling pathway in MB49 cells and reduces hypoxic fraction. A) Mechanism of activation of tarloxotinib: under severe hypoxia, tarloxotinib becomes reduced by one-electron oxidoreductases, leading to the formation of a nitro radical anion. In the absence of molecular oxygen, subsequent fragmentation of the trigger moiety in blue leads to the release of the effector tarloxotinib-E. B) MB49 cells were exposed to tarloxotinib or tarloxotinib-E for 4 hr under oxic or anoxic conditions and incubated in drug-free medium for 5 days prior to SRB staining. Values above the grouped bars are hypoxic cytotoxic ratios (oxic/anoxic IC_50_ ratio). Grouped bars represent the average IC_50_ values ± SEM from at least 4 independent experiments. C) MB49 cells were serum starved overnight and exposed to 0.1, 1 and 10 μM of tarloxotinib-E for 1 hour at 37°C including 15 min of stimulation with 50 ng/mL TGFα in the designated wells. The expression levels of phosphorylated and total proteins were analysed from 10 μg of cell lysate per condition. D) MB49 cells were injected subcutaneously into the flank of C57BL/6 mice. Tumor-bearing mice were administered with a single i.p. injection of PBS or tarloxotinib followed, after 24 hr, by i.p. injection of EF5. Tumors were harvested 90 min after EF5 injection for immunofluorescence imaging. Regions of tumor hypoxia are shown in green (EF5-Cy5) while cell nuclei are stained blue (DAPI).

Using an EGFR-dependent mouse model of bladder cancer, we report that tarloxotinib combined effectively with ICI, with improved outcomes associated with reduction in tumor hypoxia and an increased ratio of infiltrating T cells to immunosuppressive cells. Interestingly, tarloxotinib treatment was also effective against EGFR-negative tumors, an observation associated with MyD88/TLR9-dependent immunostimulatory properties that can contribute to the ICI-dependent anti-tumor response – a novel property that makes it a strong candidate to combine with ICI in the clinic.

## RESULTS

### Tarloxotinib blocks EGFR signaling and reduces the hypoxic fraction in MB49 tumors

An initial screen of syngeneic murine cancer cell lines was conducted (n=17; **Supplementary Figure 1**), identifying the transplantable murine bladder cancer cell line MB49 as sensitive to tarloxotinib **(Figure 1B)**, with an IC_50_ value of 0.62 ± 0.25 μM under aerobic (oxic) conditions, that was reduced 46-fold to 0.013 ± 0.006 μM under anoxia, demonstrating hypoxia-dependent anti-proliferative activity (*P* <0.01). In contrast, the cognate TKI, tarloxotinib-E, was active irrespective of cellular oxygenation (IC_50_ = 1.7 – 2.1 nM) **(Figure 1B)**. MB49 cells express wild-type (WT) EGFR, detected as the phosphorylated form pEGFR^Y1068^, along with its downstream signaling kinases ERK and AKT under serum-starved or standard growth conditions **(Figure 1C)**. Stimulation with transforming growth factor α (TGFα), a ligand for EGFR, enhanced levels of pEGFR^Y1068^, phosphorylated ERK (pERK1/2) and to a lesser extent phosphorylated AKT (pAKT) **(lane 3, Figure 1C)**. This response was inhibited with tarloxotinib-E without affecting total protein levels unaffected (lane 4-6). The *in vitro* sensitivity of MB49 cells was therefore associated with on-target inhibition of EGFR signaling.

Analysis of MB49 tumor lysates *ex vivo* also revealed expression of the phosphorylated proteins pEGFR, pAKT and pERK, indicating EGFR signaling was constitutively activated. A single dose of tarloxotinib at 50 mg/kg (to approximate plasma exposures observed in human subjects at 150 mg/m^2^) was sufficient to significantly reduce phosphorylation (**Supplementary Figure 2**), with inhibition of pEGFR observed within 3 hr and persisting up to 96 hr, whereas partial recovery of downstream signaling was observed after 48 hr. To examine the impact of tarloxotinib treatment on hypoxia, tumor-bearing mice were injected with EF5, a probe that binds tissues under severe hypoxia (<1 μM O_2_) and can be detected by immunofluorescence imaging (26, 27). The MB49 tumors was on average 9.56% (± 1.52%) EF5-positive and this was significantly lowered (*P* = 0.0081) 24 hr after a single dose of tarloxotinib to 1.19% (± 0.78%) **(Figure 1D)**.

### Combining immune checkpoint inhibitors with tarloxotinib improves anti-tumor activity

Next, the therapeutic potential of tarloxotinib to enhance responses to ICI was evaluated in mice with subcutaneous MB49 tumors. Administration of either anti-CTLA-4, anti-PD-1 or anti-PD-L1 monotherapies induced modest anti-tumor activity **(Figure 2A-C)**. Tarloxotinib also showed single agent activity, reflecting the impact of EGFR inhibition. Importantly, the combination of each ICI with tarloxotinib significantly potentiated activity, with a reduced rate of tumor growth and significantly improved median survival in each case (**Supplementary Figure 3A**). Notably, this was achieved without additional body weight loss (BWL) compared to the respective monotherapies (<5% average BWL) (**Supplementary Figure 3B**). In replicate experiments, combining tarloxotinib with anti-PD-L1 was consistently the most efficacious treatment; therefore, this combination was selected for further evaluation. To test causality, we sought to establish the requirement for both agents to be bioactive. No enhanced activity was observed when anti-PD-L1 was combined with a non-frangible isomer of tarloxotinib that was unable to release intratumoral tarloxotinib-E and suppress tumour hypoxia (**Supplementary Figure 4A**). Again, no enhanced activity over tarloxotinib alone was seen when combined with a non-specific IgG2b isotype antibody in place of the ICI (**Supplementary Figure 4B**). Following this initial validation we conducted a more protracted dosing schedule, involving tarloxotinib and anti-PD-L1 in three repeated cycles (with a 2-day rest between cycles), which was well tolerated (**Supplementary Figure 3C**), with extended tumor growth delay, and improved median survival (**Figure 2C)**. However, complete tumor eradication was not achieved suggesting some adaptive resistance mechanisms were still active.

**Figure 2.**
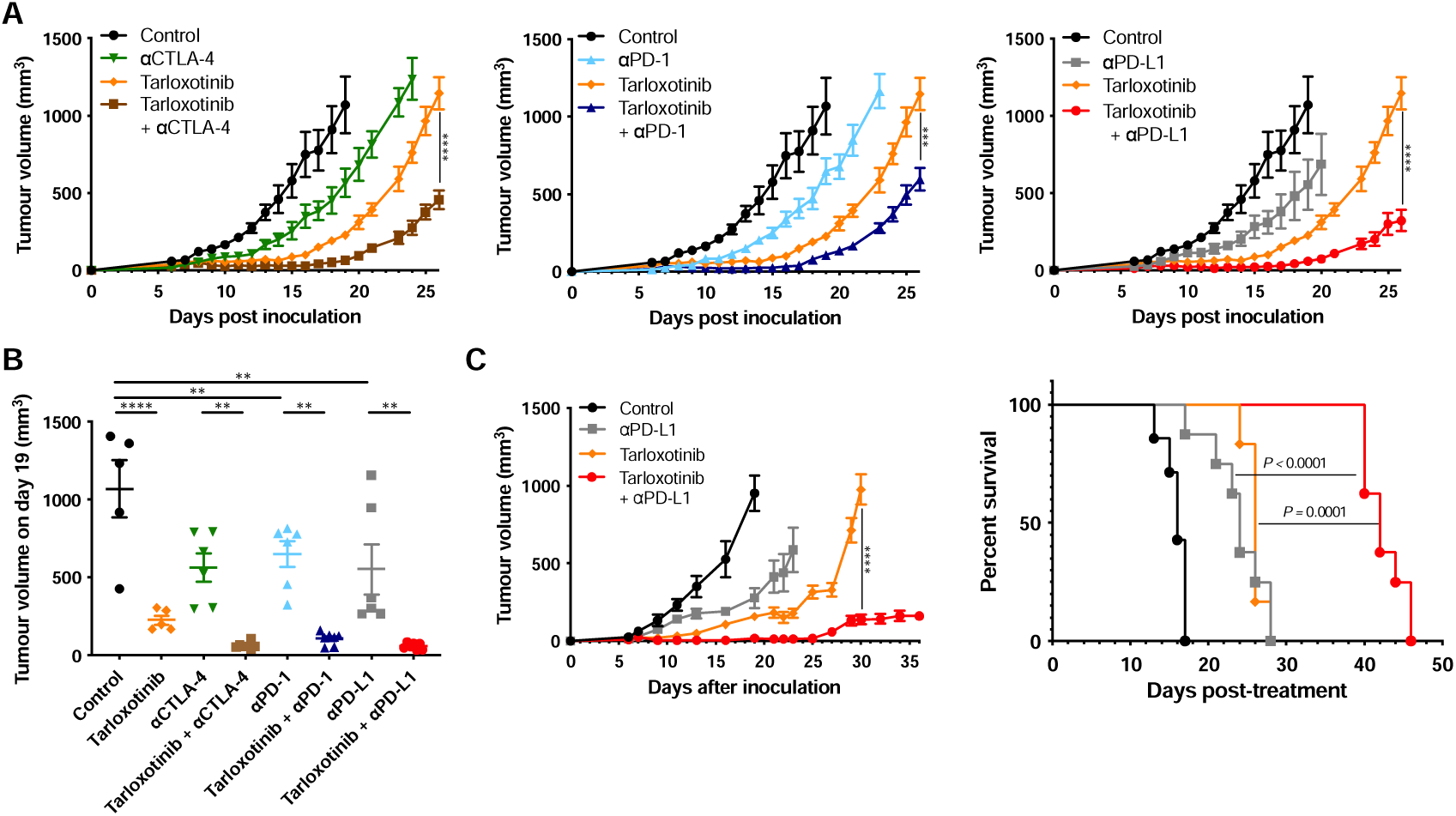
Tarloxotinib combined effectively with immune checkpoint inhibitors *in vivo* to delay MB49 tumor growth. A) MB49 cells were injected subcutaneously into the right flank of C57BL/6 mice (n = 7). From day 6, tumor-bearing mice were treated with vehicle (PBS or 20% CDW), the ICI, tarloxotinib alone or the combinations of ICI with tarloxotinib. Graphs are presented for each ICI and combination, with the same vehicle and tarloxotinib only groups serving as controls (represented in each graph for clarify). Dosing regimen for each treatment is described in Materials and Methods. Data are representative of at least two experimental repeats. B) Average volumes of MB49 tumors on day 19 post-inoculation. C) Tumor growth and survival curves (n=6-8 per treatment) after three cycles of vehicle, anti-PDL1, tarloxotinib or the combination starting from day 6. Statistical significance in tumor volumes between groups were determined using one-way ANOVA with Turkey’s multiple comparison test: **, *P*<0.01; ***, *P*<0.001; ****, *P*<0.0001. Statistical significance in survival between groups were determined using Log-rank (Mantel-Cox) test.

### Tarloxotinib alone or in combination with immune checkpoint inhibitor is associated with changes in immune profile

To identify which immune phenotypic changes were causally related to the improved anti-tumor response with the tarloxotinib/anti-PD-L1 combination, immune cell populations in the tumor and blood were examined by flow cytometry one day after treatment cessation (day 14 from tumor inoculation). The numbers of infiltrated immune cells (CD45^+^ cells) per milligram of tumor were significantly higher with the combination treatment compared to untreated controls, including significantly increased levels of CD4^+^ and CD8^+^ T cells **(Figure 3A-B)**. This included increased numbers of cytokine-producing T cells (IFNγ^+^) **(Figure 3C)**, and significantly increased proportions of CD8^+^ T cells that were proliferating (Ki67^+^) **(Supplementary Figure 5A)**. While tarloxotinib alone or in combination with anti-PD-L1 led to increased infiltration of Tregs (CD4^+^ FOXP3^+^ cells) **(Figure 3D),** the relative proportion of CD8^+^ T cells to Tregs ratio was significantly higher in the combination group compared to all other groups. The CD8/Treg ratio is considered more indicative of immune activation status than absolute numbers of Tregs (28). Expression of PD-1 on intratumoral CD4^+^ and CD8^+^ T cells **(Figure 3E)** was also enhanced with the combination treatment. In this setting, PD-1 expression is considered a marker of T cell activation and correlates with enhanced functional avidity of specific T cells (29). Treatment-induced changes in the myeloid compartment were also observed. The percentage of immune cells expressing markers typical of polymorphonuclear MDSCs (PMN-MDSC) or monocytic MDSCs (M-MDSC), discriminated on the basis of expression of Ly6G or LyC respectively, were significantly decreased with the combination treatment **(Supplementary Figure 5B)**; however, the numbers of MDSCs per milligram of tumor were not significantly different. Notably, the ratio of CD8^+^ T cells to both immunosuppressive populations was highest in the combination group. The favorable ratio of CD8/M-MDSCs was also enhanced by tarloxotinib monotherapy, indicating that the prodrug alone can have a positive impact on immune profile **(Figure 3F)**. Consistent with this interpretation, tarloxotinib alone or in combination with anti-PD-L1 also significantly increased the number of natural killer (NK), natural killer T cell (NKT) and conventional DC (cDC) cells in the tumor (**Figure 3G, Supplementary Figure 5C**).

**Figure 3.**
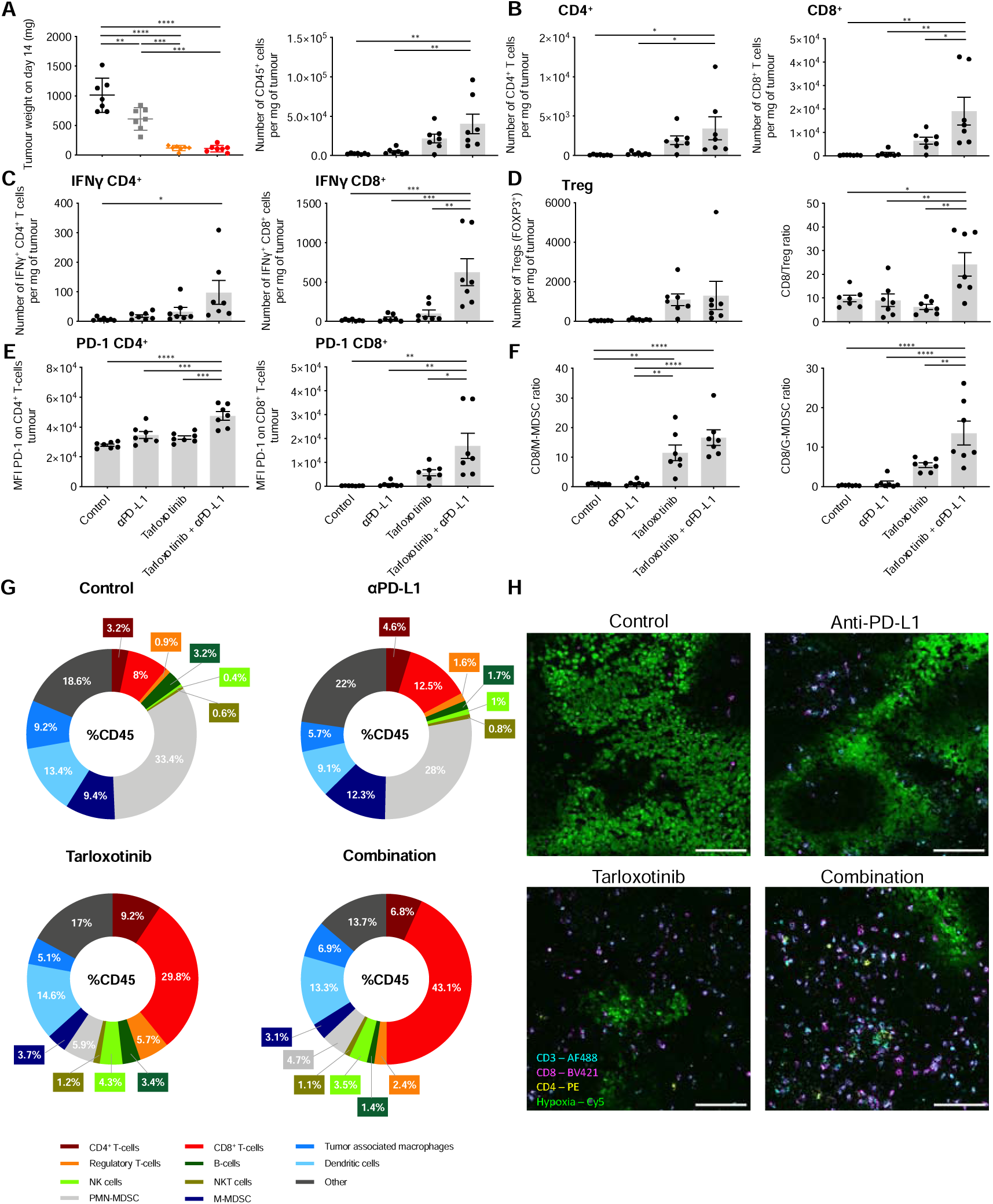
Treatment with tarloxotinib alone or in combination with anti-PD-L1 induces changes in abundance of immune infiltrates in MB49 tumors. MB49 tumor-bearing mice were treated with vehicle control, tarloxotinib, anti-PD-L1 or tarloxotinib/anti-PD-L1 combination (n = 7 per group). Tumors were harvested from mice on day 14. A-F) Average (± SEM) tumor weight, number of tumor-infiltrating immune cells, CD8/Treg and CD8/MDSC ratios and T cell PD-1 expression levels in the MB49 tumors. Relevant statistical significance between groups were determined using one-way ANOVA with Turkey’s multiple comparison test (*. *P*<0.05; **, *P*<0.01; ***, *P*<0.001; ****, *P*<0.0001). Data shown are representative of at least two experimental repeats. G) Pie-charts comparing the frequency of immune cells between untreated controls and treated MB49 tumors. H) Representative immunofluorescence images from treated MB49 tumors harvested 90 min after EF5 injection. Tumor sections were stained for CD3^+^ cells (blue), hypoxia (green), CD4^+^ cells (yellow) and CD8^+^ cells (pink). Scale bar = 100 µm.

In blood, the frequency of circulating CD4^+^ T cells was significantly higher in mice treated with the tarloxotinib/anti-PD-L1 combination than with either monotherapy (**Supplementary Figure 6A**) with comparable increase of IFNγ^+^ T cells in the peripheral blood. In contrast, the percentage of CD8^+^ T cells decreased in the blood with the combination treatment (**Supplementary Figure 6B**), although a higher proportion of these cells were producing IFNγ. This observation is consistent with migration of activated tumor-specific T cells from the blood into the tumor. Interestingly, both monotherapies led to increased expression of PD-1 on T cells relative to untreated controls, again suggesting that tarloxotinib alone can alter immune profile. However, the upregulation of PD-1 expression was more significant in mice treated with the combination in both T cell populations (**Supplementary Figure 6**).

The impact of treatment on immune profile was also assessed by immunofluorescence imaging **(Figure 3H)**. As expected, hypoxia was markedly reduced in mice treated with tarloxotinib alone or in combination with anti-PD-L1. Consistent with the flow cytometry data, control and anti-PD-L1 treated tumors showed low abundance of infiltrating CD4^+^ and CD8^+^ T cells, with imaging confirming they were largely excluded from the hypoxic zones. In contrast, administration of tarloxotinib alone resulted in infiltration of both CD4^+^ and CD8^+^ T cells into both normoxic and hypoxic tumor regions, which was significantly (*P*<0.05) enhanced when combined with anti-PD-L1 **(Supplementary Figure 7)**. Considered together, these data indicate that tarloxotinib monotherapy can have a surprisingly strong impact on the tumor-associated immune response, with more profound changes observed compared to anti-PD-L1 alone. However, the profile of these changes, including increased infiltration of T cells and higher ratio of T cells to suppressive cells, is significantly enhanced by the combination with anti-PD-L1, likely contributing to the improved antitumor responses observed.

### The improved anti-tumor response of ICI with tarloxotinib requires enhanced activity of CD8^+^ T cells

As infiltration of CD8^+^ T cells into the tumor tissue is a feature of the combination treatment, the role of these cells in the anti-tumor response was examined by using the anti-CD8α antibody to deplete hosts of CD8^+^ T cells prior to tumor challenge and initiation of treatment. The depletion strategy was effective, with no sign of re-emergence of the CD8^+^ T cells population observed in the blood 10 days after tumor challenge (**Supplementary Figure 8**). Unsurprisingly, CD8^+^ T cell depleted mice did not respond to anti-PD-L1 treatment alone, reflecting the known role of ICI in enhancing CD8^+^ T cell immunity. Interestingly, the anti-tumor activity exerted by tarloxotinib was retained in depleted mice **(Figure 4A)**, despite the change in immune profile noted earlier in these tumors **(Figure 2)**. However, the additional therapeutic benefit provided by combination with anti-PD-L1 was lost in CD8^+^ T cell depleted mice. These data suggest that CD8^+^ T cell function is specifically improved by the combined treatment, possibly because the T cells are not directly exposed to the same level of hypoxia following tarloxotinib treatment. To investigate this, EF5 was injected into tumor-bearing mice to stain CD4^+^ and CD8^+^ T cells *in situ* on day 18 after tarloxotinib was administered **(Figure 4B)**. This showed that the majority of T cells in the TME of tarloxotinib-treated mice had lower binding of EF5, and fewer cells with high EF5 intensity compared to untreated tumor-bearing controls, indicating lesser exposure to hypoxia.

**Figure 4.**
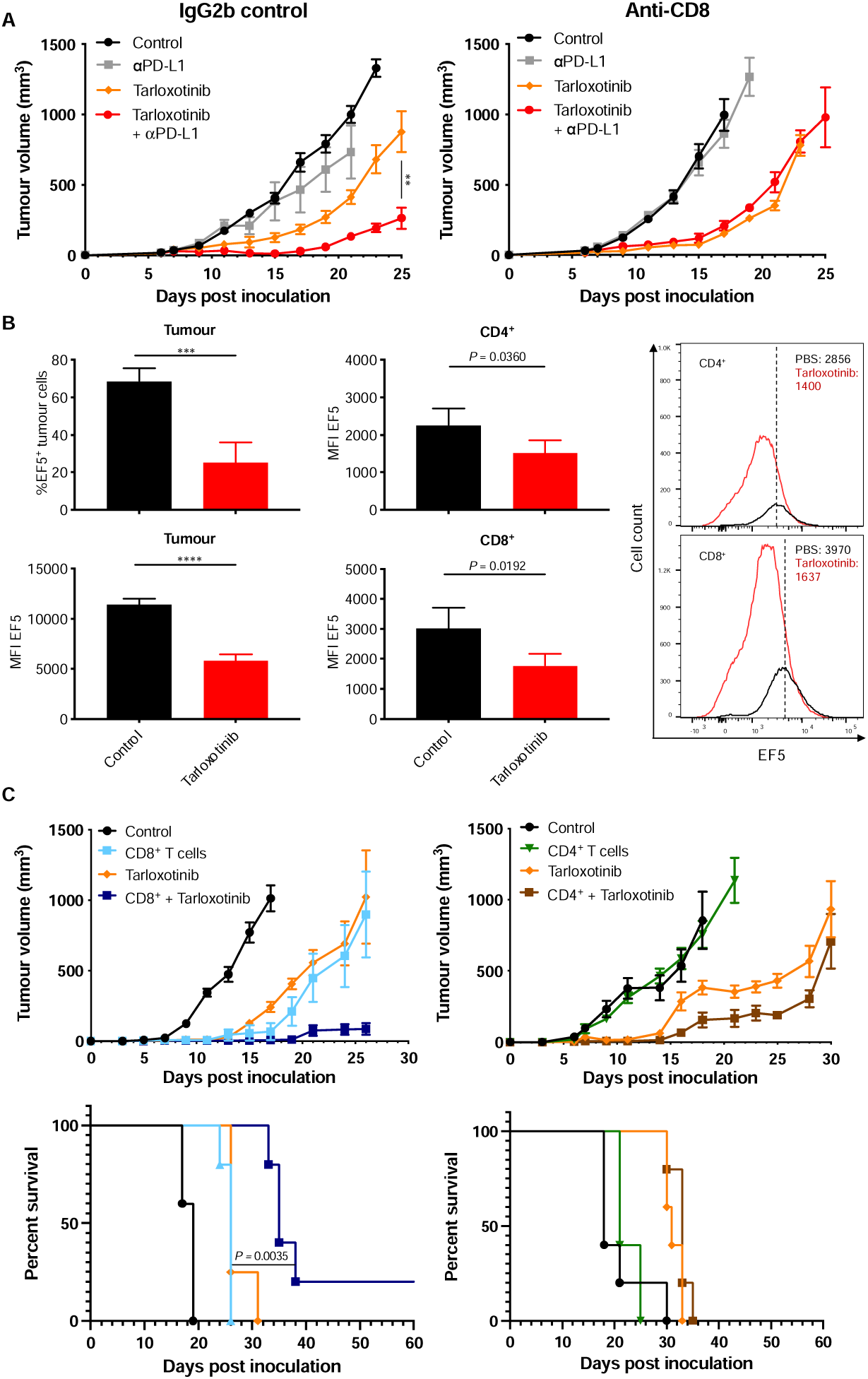
Administration of tarloxotinib reduced exposure of intratumoral T cells to hypoxia and improved the function of CD8^+^ T cells which are crucial for the combinatorial activity. A) Anti-CD8α depleting antibody or IgG2b isotype control were administered to MB49 tumor-bearing mice i.p. 2 days before tumor inoculation and 4 days before treatment initiation. Mice were randomly assigned to receive vehicle only control, αPD-L1, tarloxotinib or the combination starting from day 6. B) Left, bar graphs showing average (±SEM) (n=5 tumors per group) percentage and MFI of EF5^+^ tumor cells and MFI of EF5^+^ CD4^+^ and CD8^+^ T cells in control and tarloxotinib treated MB49 tumor-bearing mice. Right, representative flow cytometry histograms depicting differences in the number of EF5^+^ T cells between control and tarloxotinib treated tumors. Numbers represent MFI of EF5 for the respective groups and dashed lines mark the peak EF5 MFI of T cells in the control group. Statistical significance was determined using an unpaired t-test (***, *P*<0.0005; ****, *P*<0.0001). Data shown are representative of at least two experimental repeats. C) Mice were injected subcutaneously with OVA-expressing MB49 cells. Tumor-bearing mice were treated with vehicle control, tarloxotinib, CD8^+^ or CD4^+^ T cells or tarloxotinib/T cell combination (n = 5). OVA-specific T cells were activated *in vitro* by co-culturing with SIINFEKL-loaded or ISQAVHAAHAEINEAGR-loaded BMDC and then adoptively transferred to mice i.v. on day 3.

To examine the impact of tarloxotinib on T cell effector function in the absence of any confounding effect of treatment on T cell priming, an adoptive transfer model was used. The MB49 cell line was engineered to express chicken ovalbumin (OVA) so that the anti-tumor impact of *in vitro* activated OVA-specific CD8^+^ T cells (from TCR transgenic OT-I mice) or activated CD4^+^ T cells (from OT-II mice) could be examined following i.v. administration, with or without tarloxotinib co-treatment. While activated OT-I cells alone or tarloxotinib alone each demonstrated substantial anti-tumor activity, they were more effective at inhibiting tumor growth and improving survival when combined *in vivo* **(Figure 4C)**. In contrast, transfer of activated OT-II cells had no anti-tumor impact alone, and tarloxotinib treatment failed to enhance survival (**Figure 4C)**. This demonstrates that tarloxotinib can have a specific and profoundly positive impact on the function of activated CD8^+^ T cells, most likely reflecting modulation of the TME.

### Tarloxotinib demonstrates immunostimulatory activities

To further investigate the mechanism behind the observed combinatorial activity of tarloxotinib and anti-PD-L1, an mRNA transcript analysis was performed on frozen MB49 tumor extracts. Gene expression data showed tarloxotinib significantly upregulated 15 of 25 functional pathways as detected by NanoString PanCan mouse IO 360 Gene Expression Panel. Of 770 cancer/immune related genes examined, 141 were significantly upregulated and 43 downregulated (Benjamini-Yekutieli, *P*<0.05). An overview heatmap of the normalized data summaries the trends in gene expression across multiple pathways (**Figure 5A**). Most striking was the strong induction following tarloxotinib treatment of gene clusters related to co-stimulatory signaling (32 of 76 genes), immune cell adhesion and migration (32 of 75 genes), antigen presentation (14 of 52 genes), cytokine and chemokine signaling (26 of 90 genes), interferon signaling (12 of 58 genes), the lymphoid compartment (31 of 74 genes) and the myeloid compartment (17 of 69 genes). Seven of 25 pathways were also downregulated including DNA damage repair pathways, cell proliferation and the hypoxia gene cluster. Volcano plots highlight the significant differences in gene expression when the vehicle and anti-PD-L1 arms when compared tarloxotinib and combination arms (**Supplementary Figure 9)**. In contrast, gene expression in the vehicle and single agent anti-PD-L1 treatment groups did not significantly differ (*P*=0.489 for the top hit; **Supplementary Figure 9C**). This indicates tarloxotinib is the major cause of gene expression differences in these experiments.

**Figure 5.**
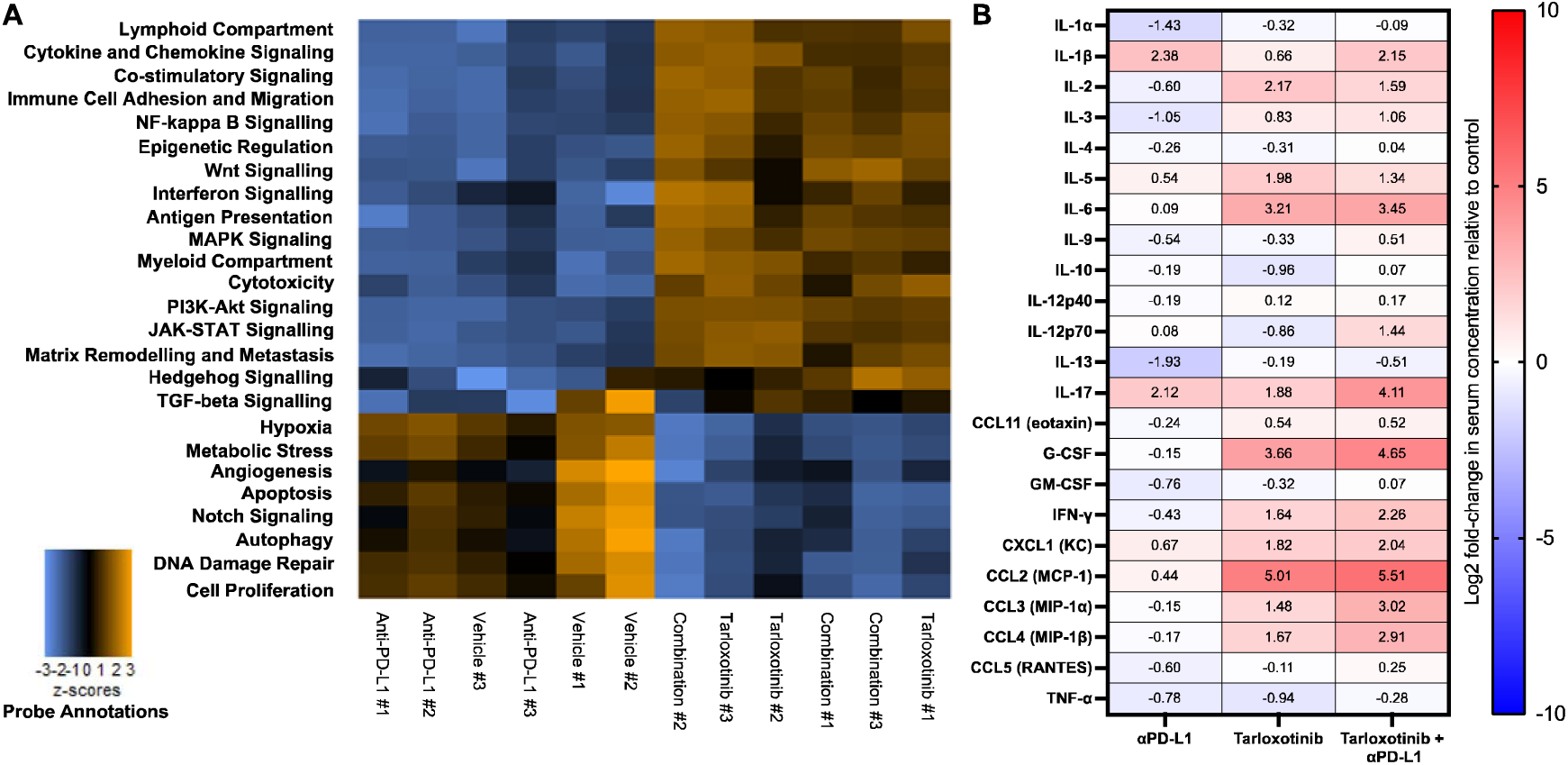
Tarloxotinib treatment resulted in amplification of multiple pathways genes in MB49 tumors and enhanced serum cytokine production. A) Overview heat-map showing relative changes in gene expression across key pathways in MB49 tumors subjected to different treatments. Rows represent different pathways and columns represent samples from different treatment cohorts. Orange color denotes gene sets with significantly upregulated expression and blue denotes gene sets with significantly downregulated expression (z-score). B) Heatmap showing the fold-changes in serum cytokine concentrations in tumour-bearing mice 6 hr after final dose with tarloxotinib, with or without anti-PD-L1, relative to untreated controls. Numbers represent the Log2 fold change after normalization of the mean concentrations to that of the control group (n=5 mice per group). Data shown are representative of two experimental repeats.

Given the gene expression profile induced with tarloxotinib included changes in chemokine and cytokine genes, we explored this further by examining protein levels in serum. Levels of several of these factors were increased in tumor-bearing mice exposed to single agent tarloxotinib, including IL-2, IL-3, IL-5, IL-6, CCL11 (eotaxin), CXCL1 (KC) and CCL2 (MCP-1), while the combination with anti-PD-L1 had no further impact **(Figure 5B)**. Tarloxotinib also induced elevation of IL-17, IFN-γ, G-CSF, CCL3 (MIP-1α) and CCL4 (MIP-1β), and serum concentrations of these factors were further enhanced when combined with anti-PD-L1 **(Supplementary Figure 10)**. In contrast, treatment with anti-PD-L1 alone did not enhance cytokine levels. Taken together, these results indicate that single agent tarloxotinib is immunomodulatory, favorably modulating immune related gene transcription and protein expression. Nonetheless, combination with anti-PDL-1 is required to exploit this effect to enhance anti-tumor responses.

### Improved anti-tumor activity of combination treatment is MyD88 and TLR9 dependent

The serum cytokine production profile associated with tarloxotinib treatment displays characteristics resembling toll-like receptor (TLR) agonism (30, 31). To test the hypothesis that tarloxotinib can stimulate TLR activation, MyD88-deficient mice were utilized as hosts for MB49 implantation. Interestingly, the single-agent activity of tarloxotinib was not influenced by the absence of MyD88 **(Figure 6A)**, yet the combination-induced improvement in anti-tumor activity typically observed in wild-type mice was absent. The diminished anti-tumor activity was also accompanied by a reduction in serum cytokine production **(Figure 6B)**. Based on evidence that tarloxotinib-E but could activate TLR9 in HEK293 TLR9 reporter cells *in vitro* (**Supplementary Figure 11**), analyses were conducted in TLR9 deficient animals. Again, the single agent activity of tarloxotinib was retained, but the combination treatment did not improve anti-tumour activity (**Figure 6C)**. These data strongly suggest that stimulation via TLR9 contributes directly or indirectly to enhance responses unleashed by anti-PD-L1.

**Figure 6.**
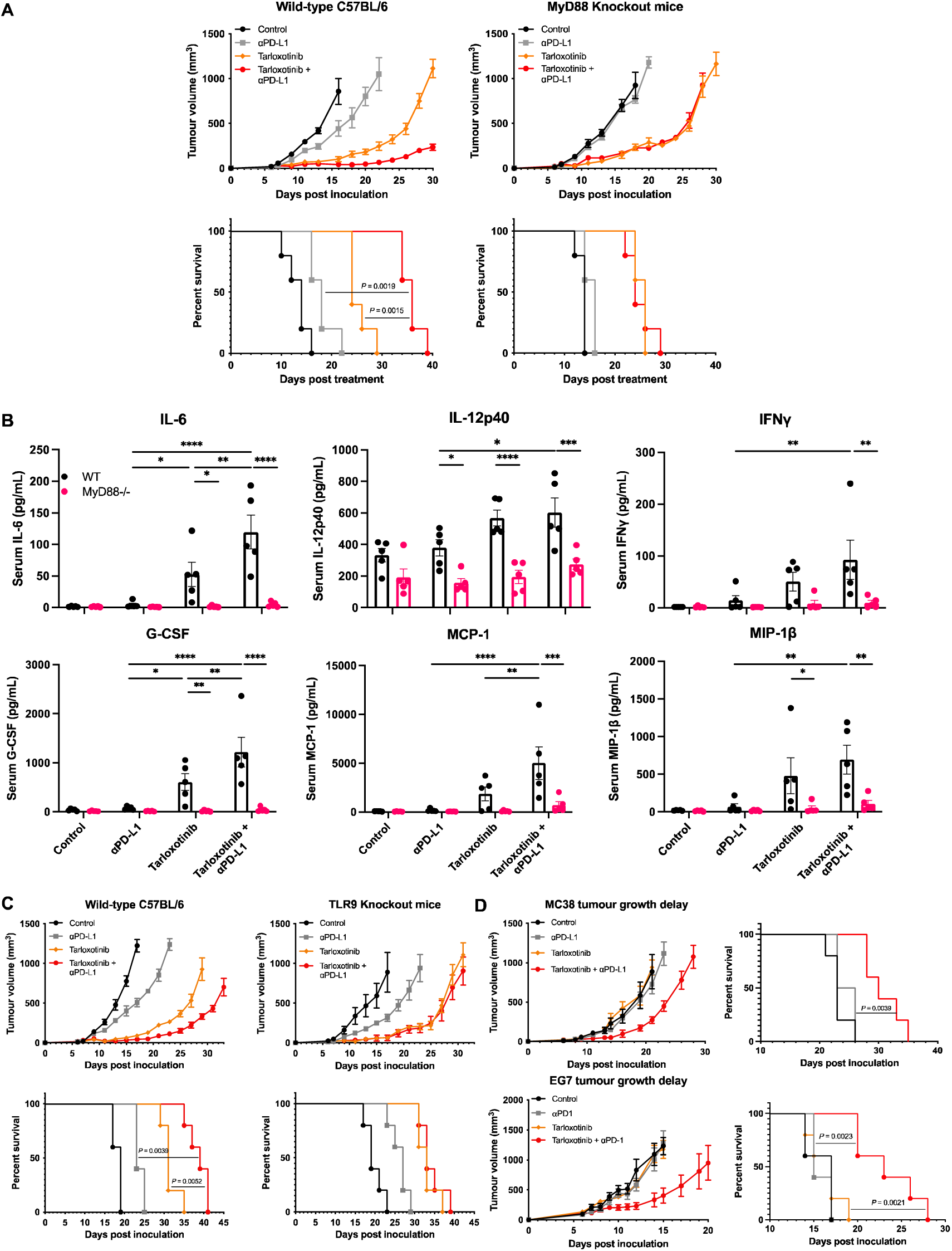
Tarloxotinib enhances anti-tumor activity and induces serum cytokine production in a MyD88-dependent manner. A) Tumor growth and survival in wild-type or MyD88 KO mice. Statistical tests conducted using Log-rank (Mantel-Cox) test. B) Serum cytokine concentrations in treated tumor-bearing WT and MyD88 KO mice. Serum was collected 6 hr post-treatment on day 14. Statistical significance between groups was determined using Two-way ANOVA with Sidak’s multiple comparison test. C) Tumor growth delay and survival in wild-type or TLR9 KO mice. Statistical significance in survival between groups were determined using Log-rank (Mantel-Cox) test. D) MC38 or EG7.OVA cells were inoculated to mice subcutaneously on the flank. Tumor-bearing mice were treated with vehicle control, tarloxotinib, anti-PD-L1/PD-1 or a combination thereof (n = 5). Data shown are representative of at least two experimental repeats.

### Potentiation of anti-tumor activity can be independent of EGFR expression

The involvement of MyD88/TLR9 signaling raises the possibility that the immunostimulatory property of tarloxotinib may be independent of EGFR inhibition within the tumor TME. To test this hypothesis, the EGFR-independent murine tumor models MC38 (colorectal cancer) and EG7.OVA (lymphoma) were employed; both cell lines are refractory to tarloxotinib-E exposure *in vitro* **(Supplementary Figure 1)**. As expected, these tumor models were correspondingly refractory to single-agent tarloxotinib treatment *in vivo* **(Figure 6D)**. Despite no single-agent ICI activity being observed in either model, a substantial tumor growth delay was evident with the combination, resulting in significantly improved median survival. This provides further evidence that tarloxotinib can combine with ICI via mechanisms other than EGFR pathway inhibition.

### Combining tarloxotinib and anti-PD-L1 does not elevate the incidence of interstitial lung disease

Critically, clinical trials combining conventional EGFR-TKI with ICI have reported an unusually high incidence of interstitial lung disease (ILD) in NSCLC patients, a serious adverse event associated with significant morbidity and mortality (32, 33, 34). We hypothesized that the tumor-targeted nature of tarloxotinib would prevent this outcome. To assess the potential for this negative interaction, lungs were harvested from MB49 tumor-bearing mice after treatment and stained with picrosirius red for quantification of collagen disposition, a surrogate marker for ILD **(Figure 7)**. In contrast to bleomycin-treated mice, which showed profound collagen deposition, as expected (35), mice treated with anti-PD-L1 or tarloxotinib alone demonstrated no evidence of increased collagen deposition relative to untreated controls and, most relevant, the combination of tarloxotinib with anti-PD-L1 displayed no increased collagen deposition.

**Figure 7.**
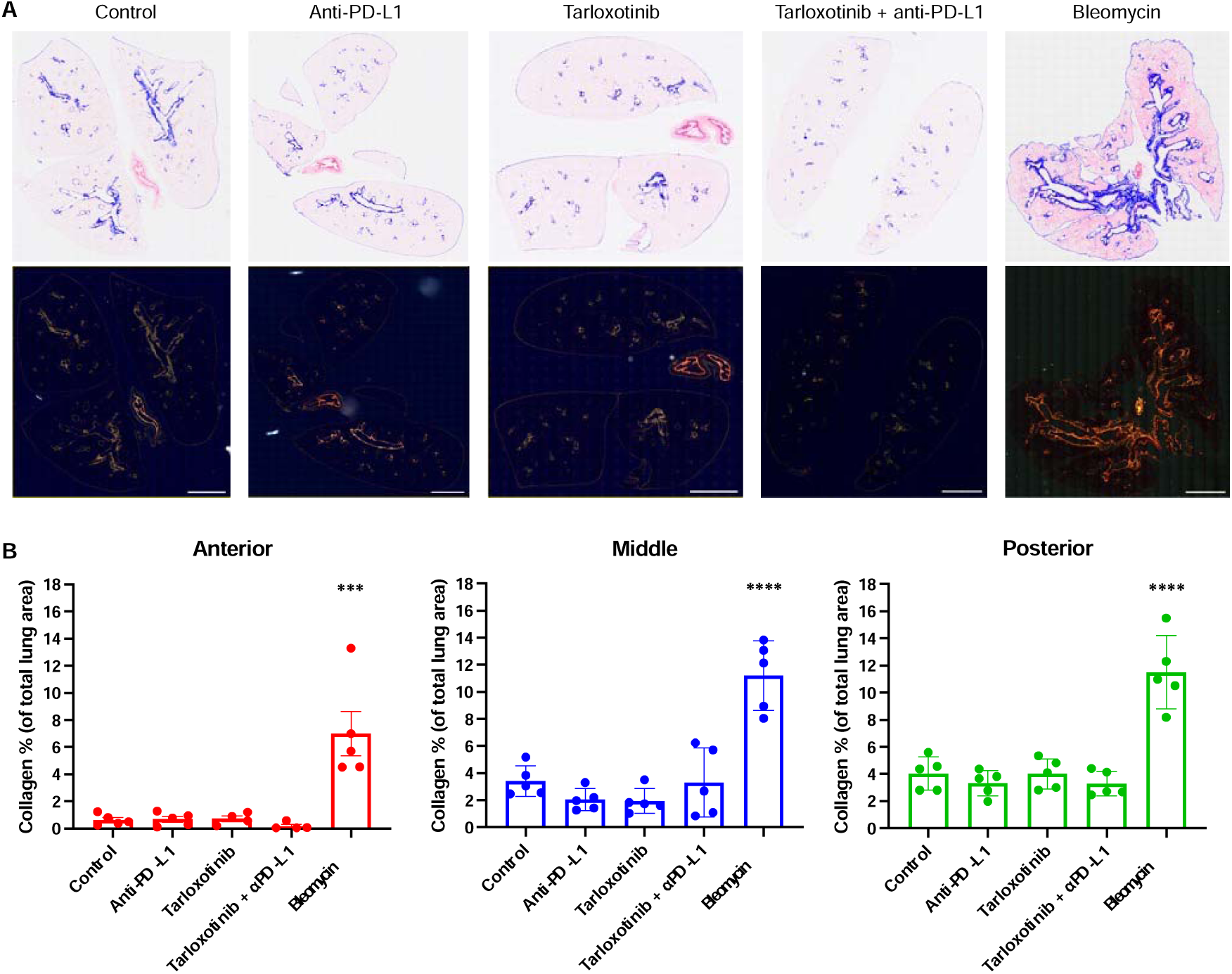
Tarloxotinib in combination with anti-PD-L1 did not induce collagen deposition in the lungs of treated mice. MB49 cells were injected subcutaneously on the flank of mice. Tumor-bearing mice were treated with bleomycin (3 mg/kg; intranasal on day 0, 2, 4 and 7), vehicle control, tarloxotinib, anti-PD-L1 or the combination thereof (n = 5). Lungs were harvested from mice on day 21, formalin fixed, sectioned and stained with picrosirius red for quantification of collagen deposition. A) Microscopy images depicting collagen staining in lungs. Top: brightfield images depicting picrosirius red staining of lung tissue with collagen positive area identified by pixel classifier shown as a blue overlay. Bottom: corresponding images in polarized view. Scale bar = 2 mm. B) Percentage of collagen of total lung area in three lung depths: anterior (50-100 µm), middle (500-600 µm from anterior) and posterior (950-1050 µm from anterior). Statistical significance between groups was determined using One-way ANOVA with Tukey’s multiple comparison test.

## DISCUSSION

There is a major clinical need for novel agents that overcome intrinsic and acquired resistance to ICI, yet very few successful adjuncts have emerged despite significant efforts (36). The efficacy of ICIs is influenced by numerous cell intrinsic and extrinsic resistance factors, with tumor hypoxia playing an important role in orchestrating suppression of the anti-tumor immune response through multiple mechanisms (3, 6). In this study, we focused on exploiting tumor hypoxia using the HAP tarloxotinib to eliminate hypoxia within the TME and thus relieve immune cells from hypoxia-mediated immune suppression. Tarloxotinib was shown to inhibit the growth of MB49 tumors through inhibition of EGFR-dependent cell signaling, whilst suppressing tumor hypoxia. Importantly, tarloxotinib combined effectively with ICIs especially with anti-PD-L1, significantly improving anti-tumor activity. Mechanistically, tarloxotinib modulates the TME, amplifying the number of infiltrating T cells, enhancing their effector function, and concurrently inducing production of pro-inflammatory cytokines in a MyD88- and TLR9-dependent fashion. Thus, tarloxotinib in combination with ICI appears to modulate the TME through independent but coordinated events, presenting a promising adjunct for overcoming ICI resistance or relapse commonly observed for many solid tumors. Critically, this tumor-targeting approach allows for increased dose intensification, without exacerbating the on-target toxicities typical of conventional EGFR-TKIs (24).

Tarloxotinib demonstrated on-target inhibition of EGFR in MB49 cells both *in vitro* and *in vivo,* effectively blocking EGFR activation and its downstream signaling through AKT and ERK. Specifically, tarloxotinib demonstrated a 46-fold increased anti-proliferative potency under hypoxia *in vitro* and *in vivo*. This was anticipated to relieve immune cells from a plethora of hypoxia-mediated immunosuppression mechanisms and induce favorable immune phenotypic changes in mice that enhances the anti-tumor immune response provided by ICIs. Indeed, profound changes in immune profile were observed in mice subjected to tarloxotinib treatment. Since infiltration of immune cells into the TME is hindered by tumor hypoxia (11, 16), the overall enhanced infiltration of CD45^+^ immune cells, especially effector CD8^+^ T cells, is consistent with the reduction in tumor hypoxia resulting from tarloxotinib treatment. Relief from hypoxia created a more favorable TME for T cell infiltration as well as enhancing the activation and function of T cells, as evidenced by their increased IFNγ expression. IFNγ is a crucial cytokine involved in generating a robust anti-tumor immune response as it enhances CD8^+^ T cell function while driving macrophage activation, enhancing antigen presentation by dendritic cells and improving cytotoxicity capacity of NK cells (37, 38). Hypoxia, mediated through HIF-1α, can suppress the production of IFNγ on T cells and reduce the frequency of IFNγ^+^T cell infiltrate in the TME (39, 40). Our observation that tarloxotinib treatment significantly reduces MB49 tumor hypoxia is consistent with the reversal of this phenomena and increased IFNγ^+^ T cell infiltration. IFNγ stimulates the immune system to promote tumor clearance, the combination treatment-induced increased frequency of IFNγ^+^ CD4^+^ and CD8^+^ T cells, and their higher serum concentration therefore collectively contributed to the enhanced anti-tumor activity. This is consistent with other studies where increased serum IFNγ and infiltration of IFNγ^+^ CD8^+^ T cells are associated with positive responses to immunotherapies (41, 42). Moreover, the reduced percentage of circulating CD8^+^ T cells in combination treated mice was consistent with net migration of circulating T cells into the primary tumor site.

The enhanced function of intratumoral CD4^+^ and CD8^+^ T cells was also attributed to their reduced exposure to hypoxia post-treatment, as evidenced by their decreased intracellular EF5 binding within the tumor. The phenomenon of enhanced T cell function following tarloxotinib treatment was not restricted to endogenous T cells. When activated OT-I cells were adoptively transferred to MB49-OVA bearing mice, subsequent tarloxotinib-induced reduction in OT-I exposure to hypoxia resulted in improved tumor growth delay and survival. However, this effect was not observed with OT-II cell transfer, indicating hypoxia has weaker impact on suppressing CD4^+^ T cell function and is consistent with the higher percentage increase in CD8^+^ T cells infiltration post-treatment. In agreement, the efficacy of tarloxotinib/ICI was negated when endogenous CD8^+^ T cells were preemptively depleted by neutralizing antibodies, again corroborating treatment-induced changes as being predominantly CD8^+^ T cell associated. Moreover, disabling either component of the combination, namely the *in vivo* release of tarloxotinib-E (through use of a non-fragmenting analogue), or the binding function of the ICI (IgG control), ablated all combinatorial activity. Collectively, these data show that the improved anti-tumor activity is a cooperative event, achieved by both reduction of tumor hypoxia to allow infiltration of more functional CD8^+^ T cells and the action of anti-PD-L1 to block the PD-1/PD-L1 inhibitory pathway to lift suppression on T cell function.

The overall number of Tregs per milligram of tumor were noticeably higher in treated mice due, in part, to the enhanced infiltration of CD45^+^ immune cells following the reduction in hypoxia. Due to negative crosstalk, the balance between the number of CD8^+^ T cells and Tregs is fundamental for a functional and effective anti-tumor immune response (43). The increased CD8/Treg ratio indicates a higher proportion of CD8^+^ T cells in the tumor avoid Treg-mediated negative regulation, enabling them to exert their cytotoxic activity. Our data are consistent with other preclinical studies in which a higher CD8/Treg ratio is associated with a more inflamed TME, leading to improved tumor clearance (44, 45, 46). The reduction in tumor hypoxia also lowered recruitment of PMN-MDSC and M-MDSC in the TME post-treatment, likely via reduced HIF-1α activity. Similar to Tregs, the CD8/MDSC ratios were higher in combination treated tumors, indicating a more selective infiltration of cytotoxic CD8^+^ T cells to the tumor. Overall, tarloxotinib modulated the TME, reducing the proportion of suppressive cells relative to CD8^+^ T cells in tumors, substantially enhancing anti-tumor immunity especially when combined with anti-PD-L1.

NanoString gene expression and serum cytokine analysis highlighted that tarloxotinib treatment possessed immunostimulatory properties which also contributed to the improved anti-tumor response observed in mice. Tumors from mice treated with tarloxotinib alone or in combination with ICI showed increased gene expression in multiple immune-related pathways. For example, genes in the interferon signaling pathway such as *Ifng, Stat1* and *Ifngr1/2* were upregulated post-treatment, consistent with the enhanced secretion of IFNγ and the increased abundance of IFNγ^+^ T cells in the blood and tumors of tarloxotinib-treated mice. The IFNγ pathway genes are crucial for responses to ICIs; indeed, tumors that are resistant to ICI commonly have genomic alterations in the IFNγ pathway (47, 48). Tarloxotinib treatment also induced production of several proinflammatory cytokines and chemokines in the serum of mice, with a profile similar to other immunostimulatory agents that activate TLRs (30, 31, 49). Of relevance, the EGFR inhibitors afatinib and osimertinib have been shown to possess immunostimulatory properties that function via MyD88, albeit at super-physiological concentrations (50, 51). MyD88 is an adaptor protein for the TLR signaling pathway, leading to AP-1 and NF-κB-dependent transcription of IL-6, IL-12 and TNF-α as well as production of type I IFNs via IRF7 (52, 53, 54). In MyD88 or TLR9 KO mice, tarloxotinib retained its single-agent activity in EGFR-dependent MB49 tumors but failed to elevate production of cytokines and chemokines. This was manifest as a lack of additional tumor growth inhibition in the anti-PD-L1 combination and is consistent with both EGFR and TLR-dependent mechanisms. To test the hypothesis these phenomena were independent, tarloxotinib was administered in the refractory, EGFR-negative MC38 and EG7 tumor models. Here, despite the absence of single-agent activity, a positive anti-tumor interaction was observed in combination with ICI, supporting the observations that tarloxotinib has immunostimulatory activities that are not dependent on EGFR-mediated effects such as downstream signal inhibition, growth inhibition and subsequent suppression of hypoxia.

The requirement for MyD88/TLR9 signaling indicates tarloxotinib may act as a dual EGFR/TLR9-targeted agent with two mechanisms of action working cooperatively. Notably, TLR9 agonists are typically large synthetic oligonucleotide molecules administered via direct intratumoral injection to enable delivery and minimize toxicity, restricting use to locally accessible solid tumors. Tarloxotinib therefore represents a clinical-stage, systemically administered small-molecule with a proven safety profile that is capable of activating TLR9 signaling in the TME to aid tumor clearance. The tumor-specific activation of tarloxotinib minimizes systemic toxicities related to EGFR inhibitors such as skin rash and diarrhea (25). Higher concentrations of tarloxotinib-E TKI can be achieved *in situ* within tumors without attendant toxicities, while also providing the necessary clinically achievable exposure. Tarloxotinib was not associated with hematological toxicities in the phase II clinical trials, but rather with manageable side effects such as low-grade rash and diarrhea (55). In our study, tarloxotinib did not have any negative impact on expansion and function of T cell and DCs, suggesting it is immune sparing. The unique properties of tarloxotinib also appear to prevent the development of ILD, a serious adverse event observed clinically with EGFR-TKI and ICI combination (32, 33, 34), further demonstrating its potential for clinical evaluation.

HAPs have been in development for decades with several showing promising clinical results (20, 22, 25). However, the challenges of clinical approval necessitate patient stratification based on tumor hypoxia status. It is conceivable that intrinsic or acquired ICI-resistance may fortuitously enrich this patient population, although prospective enrichment techniques such as CT-based radiomics are anticipated to address this clinical hurdle (56). Our study has demonstrated that tarloxotinib has significant therapeutic potential in combination with ICI, without evidence of additional or unexpected toxicities, thereby encouraging design of future clinical trials incorporating both HAP and ICI treatment modalities, particularly with incorporation of biomarker-guided identification of hypoxia to aid patient stratification (57, 58, 59).

## MATERIALS AND METHODS

### Animals and ethics statement

All animal studies were approved by the Victoria University Animal Ethics Committee and performed accordingly (AEC approval 23844, 26774 and 29650). Mice were housed and bred under pathogen-free conditions in the Biomedical Research Unit of the Malaghan Institute of Medical Research (Wellington, NZ). C57BL/6J mice were sourced from the Jackson Laboratory (Bar Harbor, ME, USA). OT-I mice expressing a transgenic mouse T cell receptor (TCR) for an H-2K^b^-binding epitope of chicken ovalbumin, OVA_257-264_ (SIINFEKL), and OT-II mice expressing transgenic TCR for the I-A^b^-binding epitope OVA_323-339_ peptide (ISQAVHAAHAEINEAGR) were originally provided by Frank Carbone, University of Melbourne, Australia; both strains were crossed in-house with the CD45.1-congenic strain B6.SJL-*Ptprc^a^Pep3^b^*/BoyJArc (Jackson Laboratory) to allow for identification of adoptively transferred transgenic cells in CD45.2 recipients. TLR9-deficient C57BL/6J-*Tlr9^M7Btlr^*/Mmjax mice and MyD88-deficient B6.129P2(SJL)-*Myd88^tm1.1Defr^*/J mice were obtained from the Jackson Laboratory.

### Cell culture

Cell lines were maintained in a humidified incubator (37°C, 5% CO_2_) in media supplemented with 10% fetal bovine serum (FBS) and 1% penicillin-streptomycin (P/S). Authenticated MB49 cells were provided by Dr. Michael Curran (MD Anderson, TX, USA). MC38 and EG7.OVA cell lines were purchased from American Type Culture Collection (USA) and all transfected cell lines were generated at the ACSRC, University of Auckland. MB49 and MC38 cells were cultured in Dulbecco’s modified eagle medium (DMEM). EG7.OVA cells were cultured in Iscove’s Modified Dulbecco’s medium (IMDM). All culture media and supplements were purchased from Thermo Fisher Scientific (Carlsbad, CA, USA).

### *In vitro* analysis of EGFR activity

MB49 tumor cells were cultured in αMEM media containing 10% FBS, 1% P/S, 1% GlutaMAX (Gibco, CA, USA) and seeded into 96-well plates at 2000 cells/well. Plates were then transferred into either a humidified incubator (aerobic) or Bactron Anoxic Chamber (Sheldon Manufacturing Inc., OR, USA) to obtain palladium/H_2_ catalyst-induced anoxia (<1 ppm O_2_) and incubated for 2 hr to allow cell attachment. All materials were anoxia equilibrated for 72 hr prior. Following 4 hr exposure to tarloxotinib or tarloxotinib-E, cells were washed with fresh DMEM, and cultured drug-free for 5 days before staining with 0.4% Sulforhodamine B (Sigma-Aldrich, MO, USA) to determine cell density by measuring absorbance at 490 nm using the EL808 Absorbance Microplate Reader (Bio-Tek instruments, VT, USA). IC_50_ values were determined using KC4 microplate data analysis software (KC4, Bio-Tek), calculated as the interpolated drug concentration required to reduce absorbance to 50% of controls on the same plate.

### Western blotting

Cells were lysed in RIPA lysis buffer (50 mM/L Tris-HCl, 1% nonidet P40 substitute, 0.25% sodium deoxycholate, 150 mM/L NaCl and 1 mM/L EDTA) containing 1:100 protease inhibitor (Sigma-Aldrich, MO, USA). Cell lysates were incubated on ice for 30 min and the non-soluble components were separated from supernatants containing proteins by centrifugation at 13,000 rpm (5 min, 4°C). Tumors were harvested from mice and snap frozen in liquid nitrogen, then cryofractured and homogenised with Laemmli buffer containing 2% sodium dodecyl sulphate, 0.125 M Tris-HCl, 10% glycerol, 5% β-mercaptoethanol and protease inhibitor (1:1000). After vortexing, samples were heated to 70°C with agitation at 300 rpm for 10 min, and undissolved tissues were pelleted and removed by centrifugation at 13,000 rpm (5 min, 4°C). For phosphorylated proteins, Laemmli buffer was supplemented with 1 mM phosphatase inhibitors sodium orthovanadate and sodium fluoride. Samples were subjected to SDS-PAGE electrophoresis (100V, 90 min) and transferred to nitrocellulose membranes (Bio-Rad, CA, USA) by electrophoresis (100V, 1 hr) in a pre-assembled transfer cassette in a chamber containing cold transfer buffer (25 mM Tris, 200 mM glycine and 20% methanol). Following transfer, membrane was rinsed in 5% PhosphoBlocker Blocking Reagent (Cell Biolabs Inc., CA, USA) in TBS-T (tris-buffered saline with 1% Tween-20) and incubated overnight at 4°C with primary antibodies at concentrations in **Supplementary Table 1**. The membranes were then washed with TBS-T and incubated with secondary horseradish peroxidase-conjugated antibodies (Santa Cruz Biotechnology Inc., CA, USA) (2 hr, RT). Protein bands were detected by chemiluminescence and imaged using the ImageQuant LAS-4000 Imager (GE Healthcare, UK).

### Tumor implantation and tumor growth delay study

Mice were injected subcutaneously with 1.5×10^5^ MB49 cells, 6×10^5^ MC38 cells or 5×10^5^ EG7.OVA cells in sterile PBS. Tumor growth was monitored every 1-3 days, and volume was calculated as π × (length × width × width)/6. Six days after inoculation, mice were randomly assigned to treatment groups (5-7 mice per group). Each cycle of treatment consists of: tarloxotinib administration via intraperitoneal injection (i.p.) at 50 mg/kg every 2 days for 4 doses total (q2dx4); immune checkpoint inhibitors (all BioXCell, NH, USA) including anti-PD-1 (clone RMP1-14), anti-PD-L1 (clone 10F.9G2) and anti-CTLA-4 (clone UC10-4F10-11) administration i.p. at 200 µg per mouse every 3 days for 3 doses total (q3dx3). For *in vivo* depletion of T cells, 200 µg of anti-CD8α (clone 2.43) or anti-CD4 (clone GK1.5), from BioXCell (NH, USA), were administered i.p. 2 days before tumor inoculation and 4 days before treatment initiation, with depletion of target population confirmed by flow cytometry analysis of peripheral blood. Relevant isotype controls (clones LTF-2 and 2A3) were administered i.p. following the same dosing schedule. Mice were euthanized when tumor volume exceeded 1300 mm^3^, when body weight loss exceeded 15% of the pre-treatment weight, extensive tumor ulceration occurred or when signs of severe adverse drug toxicities were observed.

### Immunofluorescence imaging

Tumor-bearing mice were administered the hypoxia probe EF5 (Merck, Germany) formulated in PBS, injected i.p. at 60 mg/kg. Tumors were excised 90 min later and embedded in cassettes with Tissue-Tek OCT compound (Sakura, USA). Samples were snap-frozen, sectioned (10 µm), mounted on poly-L-lysine-coated slides and fixed in cold acetone for 5 min before blocking with StartingBlock T20 blocking buffer (Thermo Fisher Scientific, USA) for 1 hr at RT. Post-wash, slides were stained with anti-EF5 Cy5 or Cy3-conjugated antibody (1 hr, RT) for detection of tumor hypoxia. For immune cells, fluorophore-conjugated antibodies against cell surface markers were used at concentrations in **Supplementary Table 2**. Slides were stained with DAPI antibody at 1:2000 for 10 min, washed and mounted in ProLong Gold Antifade Mountant (Thermo Fisher Scientific, USA), and imaged using Olympus VS200 Slide Scanner. To determine hypoxic fraction, total viable area and hypoxia positive areas were quantified.

### Multiparameter spectral flow cytometry analysis

Cell suspension samples were generated from tumor, spleen and lymph nodes harvested from tumor-bearing mice. Samples were washed in PBS and incubated with Zombie NIR Fixable Viability dye (1:1000, BioLegend) for 15 min at RT. Non-specific Fc receptor-mediated antibody binding was blocked by incubation with Fc-block (clone 2.4G2) (10 min, 4°C). Washed cells were then stained with fluorophore-conjugated antibodies (20 min, 4°C) against cell surface antigens in flow buffer containing 1:5 Brilliant Buffer Plus (BD Biosciences). For intracellular staining, cells were fixed and permeabilised using the Transcription Factor Buffer Set (BD Biosciences) by following the manufacturer’s protocol. Flow sample acquisition was performed on an Aurora (Cytek Biosciences) flow cytometer and analysed using FlowJo software (v10.6, TreeStar, OR, USA). Mode detailed methods with antibody dilution, target, source and clones are in **Supplementary Table 3.**

### Multiplex analysis of serum cytokines

Serum was collected from mice 6 and 24 hr post-treatment to measure mouse cytokine levels by Bio-Plex multiplex immunoassays (Bio-Rad Laboratories, CA, USA) following the manufacturer’s protocol. Plates were washed using the Bio-Plex Pro wash station (Bio-Rad Laboratories, CA, USA) with a magnetic plate holder. Fluorescence was recorded using the Luminex 200 array reader (Merck Millipore, MA, USA) and cytokine concentrations were determined against commercial standards (Bio-Rad Laboratories, CA, USA) using Luminex xPONENT (Luminex Corporation, TX, USA).

### *In vitro* activation and adoptive transfer of OT-I cells

To prepare activated OT-I cells, lymph nodes were harvested from OT-I mice and processed to single cell suspension. To stimulate activation, murine bone marrow derived dendritic cells (BMDC) were prepared. Briefly, single cell suspensions were prepared from bone marrow extracted from mouse femurs, and then cultured (37°C/5% CO_2_) in 6-well plates at 2×10^6^ cells per well in IMDM media supplemented with 5% FBS and 200 µl GM-CSF (PeproTech Inc., NJ, USA), with partial media renewal every 2 days. On day 7, BMDCs were matured with 200 ng/mL of LPS added overnight then washed and resuspended at 1 x 10^6^ cells/mL before incubation with 0.1 µM OVA_257-264_ peptide (4 hr, 37°C) (Genscript Biotech, NJ, USA). Free peptides were removed by washing cells in IMDM. To initiate T cell activation, 5 x 10^5^ OT-I cells were added to wells containing 6.25 x 10^4^ mature BMDCs in complete IMDM containing IL-2 (100 U/mL) and IL-7 (6.2 ng/mL) (PeproTech Inc., NJ, USA). After three days cells were ready for adoptive transfer, with 2 x 10^6^ cells resuspended in 200 µL PBS injected via the lateral tail vein.

### RNA isolation

Tumors were harvested from and snap frozen in OCT compound (Sakura Finetek, USA). Frozen tumors were cut into 10 µm sections and total RNA extracted using a Quick-RNA Microprep Kit (Zymo Research, CA, USA). In brief, tumor sections were lysed, homogenised in 300 μL RNA lysis buffer, centrifuged (>12,000 *g,* 1 min at RT) and transferred to RNase-free tubes. Samples were mixed with 300 μL of 100% ethanol and transferred into Zymo-Spin IC columns in collection tubes. After centrifugation, samples were incubated with DNase Reaction Mix I (15 min, RT) and washed with RNA wash buffer followed by RNA Prep Buffer plus two repeats of washing. DNase/RNase-free water was added to elute the concentrated RNA, which was stored at −80°C. Gene expression analysis was conducted using the PanCancer Mouse IO 360 Gene Expression Panel on a nCounter FLEX Analysis System (NanoString Technologies, WA, USA). Raw gene expression data was analysed using the Advanced Analysis Module version 2.0.115 of nSolver Analysis software version 4.0 (NanoString Technologies, WA, USA).

### Statistical analysis

Statistical analysis was performed using Prism version 10 software. Unless otherwise specified, student’s unpaired, two-tailed t-test was used to calculate statistical significance between two groups of data. One-way analysis of variance (ANOVA) with Tukey’s post-test was used to determine statistical significance between multiple groups. Two-way ANOVA with Šídák’s multiple comparisons test was used to determine statistical significance between multiple groups in presence of two categorical independent variables. Kaplan-Meier curves were used to model survival, and significance between groups were determined using Log-rank (Mantel-Cox) test. *P* values less than 0.05 were considered to be statistically significant.

