## Supplementary Method for "Tarloxotinib targets tumor hypoxia to improve therapeutic efficacy of immune checkpoint inhibition in a TLR9 dependent manner"

***SUPPLEMENTARY METHODS***

**Tissue processing for flow cytometry and cytokine analysis**

Lymph nodes, spleens and tumours were collected on day 14 after tumour inoculation and digested in media containing 100 µg/mL Liberase II/DNase (Roche, Germany) for 20 min at 37°C and then filtered through 70 μm cell strainers. Single cell suspensions were washed, centrifuged (1600 rpm, 4 min) and resuspended in RBC lysis buffer (Qiagen, USA) for 2 min. Cells were then centrifuged (2000 rpm, 2 min), washed and resuspended in flow buffer (PBS containing 1% FBS, 0.01% NaN_3_ and 2mM EDTA) for antibody staining. Peripheral blood was collected via submandibular bleed into tubes containing 200 μL of 10 mM EDTA-PBS. Red blood cells were then lysed with RBC lysis buffer (20 min, 37°C) and the remaining white blood cells were resuspended in 200 μL flow buffer for antibody staining. For serum collection, peripheral blood was collected in Microvette 500 Z-Gel blood collection tubes (Kent Scientific, CT, USA), centrifuged at 10,000 rpm for 5 min at RT and the resulting serum was transferred into 96-well plates (Nunc, USA) for storage at -20°C until required.

**Immunofluorescence imaging for quantification of T cells**

Tumour-bearing mice were treated and after 24 hr, the hypoxia probe EF5 (formulated in PBS) were administered to mice i.p. at 60 mg/kg. Tumours were excised 90 min post-EF5 injection and embedded in moulds with Tissue-Tek O.C.T compound. Samples were processed following methods adapted from Schmidt *et al.,* (2018). Briefly, 8 µm tumour cryosections were fixed in 4% paraformaldehyde (PFA) for 5 minutes before incubation with blocking buffer (10 mg/mL BSA and 1 µg/mL 2.4G2 Fc Block diluted in SuperBlock Blocking Buffer) for 1 hr at 37°C. Slides were stained with anti-EF5 Cy5-conjugated antibody (1 hr, RT) for detection of tumour hypoxia. For immune cell staining, sections were incubated with anti-CD3-AF488, anti-CD4-PE, and anti-CD8-BV421 antibodies, and the nucleic acid stain SYTOX™️ Blue overnight. Samples were washed, and mounted with Faramount aqueous mounting medium. An IX83 Inverted Microscope (Olympus, Japan) with a FV3000 Laser Scanning Confocal unit (Olympus, Japan) was used to generate four 1.997 mm^2^ stitched images at 20 × magnification (UPLSAPO 20, 0.75 NA, Olympus, Japan) within each tumour section. Images were acquired as Z-stacks with a pixel size of 0.663 µm^2^. BV421, AF488, PE, and Cy5 fluorophores were excited with 405 nm (50 mW), 488 nm (20 mW), 561 nm (20 mW), and 647 nm (40 mW) lasers, respectively. Emission of these fluorophores was detected at 423/13 nm, 520/20 nm, 595/25 nm, and 700/50 nm, respectively. In a secondary phase, SYTOX™️ Blue was excited with a 445 nm laser (75 mW), and emission was detected at 465/15 nm. FluoView V2.4.1 software was used for image acquisition. For image analysis, compensation of SYTOX™️ Blue spectral overlap with PE was performed using the ImageJ/FIJI V2.14.0 (Schindelin *et al.,* 2012) Image Calculator to subtract nuclear spill-over. Maximum intensity Z-projections were generated, and images were and imported into QuPath V0.5.1 (Bankhead *et al.,* 2017). Cells were segmented based on cell nuclei SYTOX™️ Blue staining, and a training image was generated from a randomly selected 500 µm^2^ region from 25% of the experimental images. Using the training image, random trees (Rtrees) based object classification models were trained to detect positive CD3, CD4, and CD8 staining. For hypoxia detection, a threshold-based pixel classifier was trained to detect Cy5 positive area. The optimised classifiers were applied to all images. For immune cells, total and hypoxic CD3^+^, CD3^+^CD4^+^CD8^-^, and CD3^+^CD8^+^CD4^-^ cells within the four images were quantified as a percentage of all cells. For hypoxic fraction (HF), total viable area of each section and hypoxia positive areas were quantified. At least 5 tumours per treatment group were analysed to calculate the average intratumoral T cell infiltration.

**Assessment of TLR9 activity in vitro**

To assess TLR9 activity, a range of concentrations of tarloxotinib-E or afatinib were incubated in 96-well plates with mouse (5 x 10^4^ cells/well) or human (8 x 10^4^ cells/well) TLR9 HEK-Blue SEAP reporter cells (InvivoGen, CA, USA). The TLR9 agonist ODN1826 and ODN2006 served as positive controls for mouse and human cells, respectively. The cells were cultured for 18 h in HEK-Blue detection media at 37°C in 5% CO_2_ for 16 hr. Absorbance at 620 nm was determined as a readout of TLR9-induced SEAP activity, determined using an Infinite M1000 Pro plate reader (Tecan, Switzerland).

**Quantification of collagen deposition**

Coronal sections of FFPE mouse lungs (4 µm) were stained with Picrosirius red (PSR) connective tissue stain following the manufacturer protocol, allowing for visualisation of type I and III collagen. Briefly, lung sections were dewaxed, rehydrated and stained with PSR for 1 hr at RT. After washing with water, sections were dehydrated and mounted in DPX. Whole-section imaging was performed using both brightfield and circular polarized light (45° stage rotation) imaging techniques within the VS200 Slide Scanner (Evident, Japan) fit with the IRCA-Flash4.0 C13440 Digital Camera (Hamamatsu, Japan). Images were acquired at 20 × magnification (Olympus, UPLAXAPO 20, 0.80 NA) with a pixel size of 0.2738 µm^2^. For analysis, images obtained by brightfield imaging were used to generate a whole-tissue annotation, and the annotation was copied to the corresponding polarised image. A training image consisting of randomly selected 2 mm^2^ regions from all images was used to objectively train a collagen threshold classifier set to 2.19 µm/pixel resolution. The saved classifier was applied to all polarised images. The total tissue area (mm^2^) and collagen area (mm^2^) measurements were exported, and collagen as a percentage of total section area was calculated. Results (mean ± SD) were analysed via two-way repeated measures or one-way ANOVA, followed by a Tukey’s multiple comparison test when significance was identified. All analysis was performed in Prism 10.0 with n=5 lungs per treatment group.

**Supplementary Table 1.** Antibodies used for western blotting.

| **Target protein** | **Antibody type** | **Source** | **Dilution** |
| --- | --- | --- | --- |
| Actin | Mouse monoclonal | Merck, USA | 1:10000 |
| EGFR | Rabbit polyclonal | Cell Signaling Technology, USA | 1:5000 |
| Phosphorylated EGFR (Y1068) | Rabbit polyclonal | Cell Signaling Technology, USA | 1:2000 |
| Akt | Rabbit polyclonal | Cell Signaling Technology, USA | 1:2000 |
| Phosphorylated Akt (S473) | Rabbit polyclonal | Cell Signaling Technology, USA | 1:2000 |
| Erk1/2 (p44/42 MAPK) | Rabbit polyclonal | Cell Signaling Technology, USA | 1:2000 |
| Phosphorylated Erk1/2 (T202/Y204) | Rabbit polyclonal | Cell Signaling Technology, USA | 1:2000 |
| Secondary | Goat anti-rabbit IgG-HRP | Santa Cruz Biotechnology Inc, CA, USA | 1:5000 |

**Supplementary Table 2.** Antibodies used for multicolour immunofluorescent imaging. Marker, fluorophore, clone, identifier, source and dilution are indicated.

| **Marker** | **Fluorophore** | **Identifier (clone, RRID)** | **Source** | **Dilution** |
| --- | --- | --- | --- | --- |
| CD3e | AF488 | 500A2, AB_253658 | Invitrogen | 1:400 |
| CD4 | PE | GK1.5, AB_312693 | BioLegend | 1:400 |
| CD8α | BV421 | 53-6.7, AB_273847 | BD Horizon | 1:200 |
| EF5 | Cy5 | Clone ELK3-51 | Millipore | 75 µg/mL |
| Nuclei | SYTOX^TM^ Blue | N/A | Invitrogen | 10 µM |

**Supplementary Table 3.** Antibody panel used for spectral flow cytometry experiment. Marker, fluorophore, clone, identifier, source and dilution are indicated.

| **Marker** | **Fluorophore** | **Identifier (clone, RRID)** | **Source** | **Dilution** |
| --- | --- | --- | --- | --- |
| B220 | FITC | Clone RA3-6B2, AB_312991 | BioLegend | 1:600 |
| CD3e | BV510 | Clone 145-2C11, AB_2565879 | BioLegend | 1:100 |
| CD4 | PerCP eFluor 710 | Clone RM4-5, AB_1834431 | eBioscience | 1:800 |
| CD8a | AF700 | Clone 53-6.7, AB_493703 | BioLegend | 1:600 |
| CD11b | BB700 | Clone M1/70, AB_2744272 | BD Biosciences | 1:800 |
| CD11c | BV786 | Clone HL3, AB_2738394 | BD Biosciences | 1:300 |
| CD44 | PE-Cy5 | Clone IM7, AB_468749 | eBioscience | 1:800 |
| CD45.2 | APC-Fire 750 | Clone 104, AB_2629723 | BioLegend | 1:600 |
| CD69 | SB600 | Clone H1.2F3, AB_2688097 | eBioscience | 1:600 |
| CD274 (PD-L1) | PE-Dazzle 594 | Clone 10F.9G2, AB_2565639 | BioLegend | 1:600 |
| CD279 (PD-1) | PE-Cy7 | Clone 29F.1A12, AB_10696422 | BioLegend | 1:500 |
| F4/80 | AF647 | Clone BM8, AB_893492 | BioLegend | 1:600 |
| FOXP3 | AF532 | Clone FJK-16s, AB_11218870 | eBioscience | 1:100 |
| IFNγ | PE | Clone XMG1.2, AB_395376 | BD Biosciences | 1:100 |
| Ki67 | BV480 | Clone B56, AB_2739511 | BD Biosciences | 1:200 |
| KLRG1 | APC | Clone 2F1/KLRG1, AB_10641560 | BioLegend | 1:900 |
| Ly6C | BV570 | Clone HK1.4, AB_10896061 | BioLegend | 1:500 |
| Ly6G | BV711 | Clone 1A8, AB_2738520 | BD Biosciences | 1:600 |
| MHC-II (IA-IE) | eFluor 450 | Clone M5/114.15.2, AB_1272204 | eBioscience | 1:900 |
| NK1.1 | BV650 | Clone PK136, AB_2738617 | BD Biosciences | 1:500 |
| TNFα | BV750 | Clone MP6-XT22, AB_2801090 | BioLegend | 1:400 |
| OVA pentamer | PE | OVA_257-264_ peptide-loaded H-2K^b^ pentamers | NIH | 0.5 µl/sample |
| Viability | NIR Zombie | N/A | BioLegend | 1:1000 |

**Supplementary Table 4**. Antibodies used for conventional flow cytometry experiment. Marker, fluorophore, clone, identifier, source and dilution are indicated.

| **Marker** | **Fluorophore** | **Identifier (clone, RRID)** | **Source** | **Dilution** |
| --- | --- | --- | --- | --- |
| CD3 | FITC | Clone 145-2C11, AB_312671 | BioLegend | 1:100 |
| CD4 | PerCP eFluor 710 | Clone RM4-5, AB_1834431 | eBioscience | 1:800 |
| CD8 | BUV395 | Clone 53-6.7, AB_2732919_ | BD Biosciences | 1:800 |
| CD45.2 | APC-Fire 750 | Clone 104, AB_2629723 | BioLegend | 1:600 |
| EF5 | Cy5 | Clone ELK3-51 | Millipore | 75 µg/mL |
| Nuclei | NIR Zombie | N/A | BioLegend | 1:1000 |

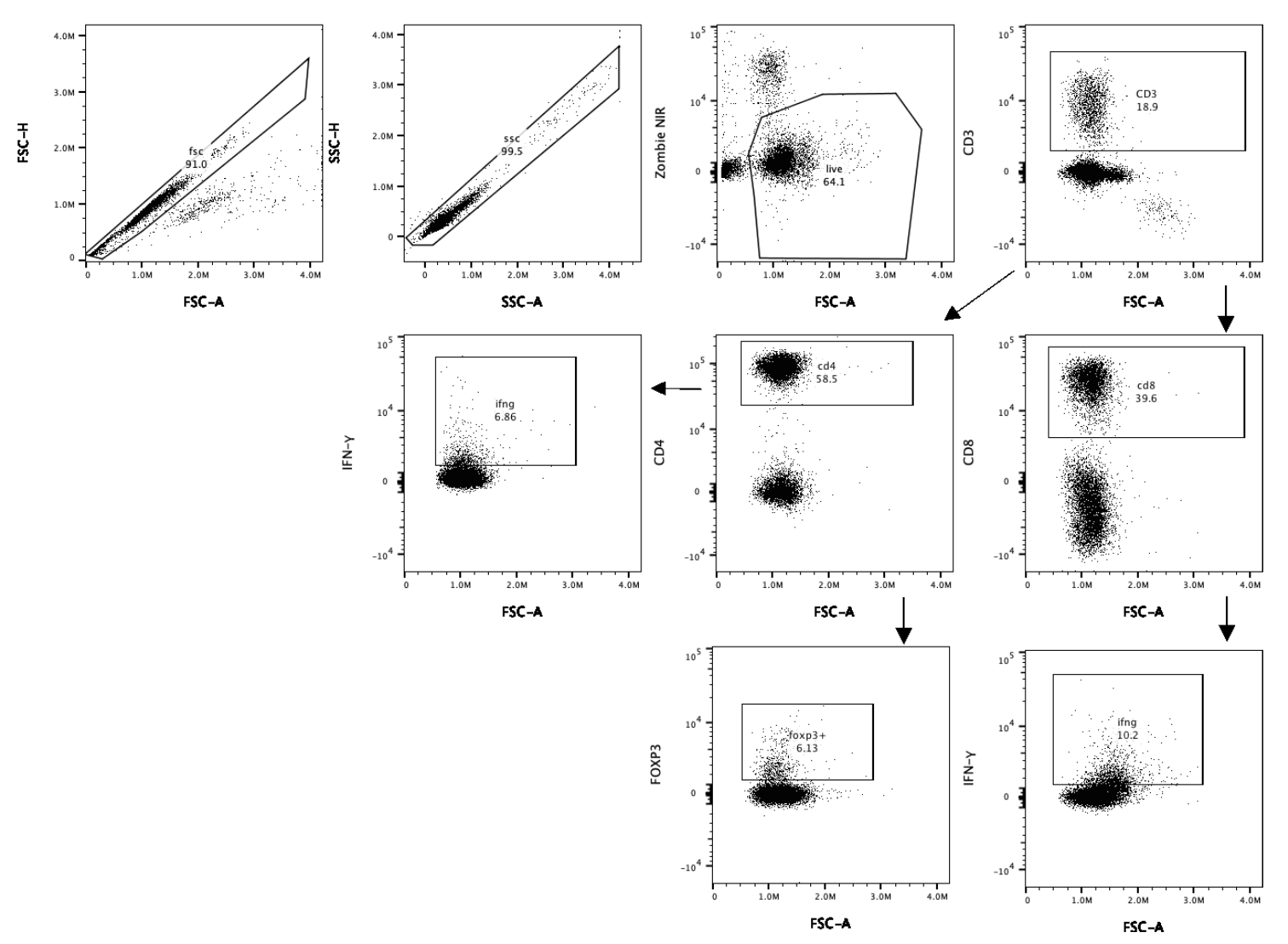

**Gating strategy of flow cytometry analysis of T cell populations.** The gating example shown is on tumour cells from treated mice. After gating on singlets and cells of interest (the gate was set in the lymphocytes region using the scatter profile of the cells), leukocytes were defined by staining with anti-CD45, and then the strategy shown was used to identify various T cells subtypes.

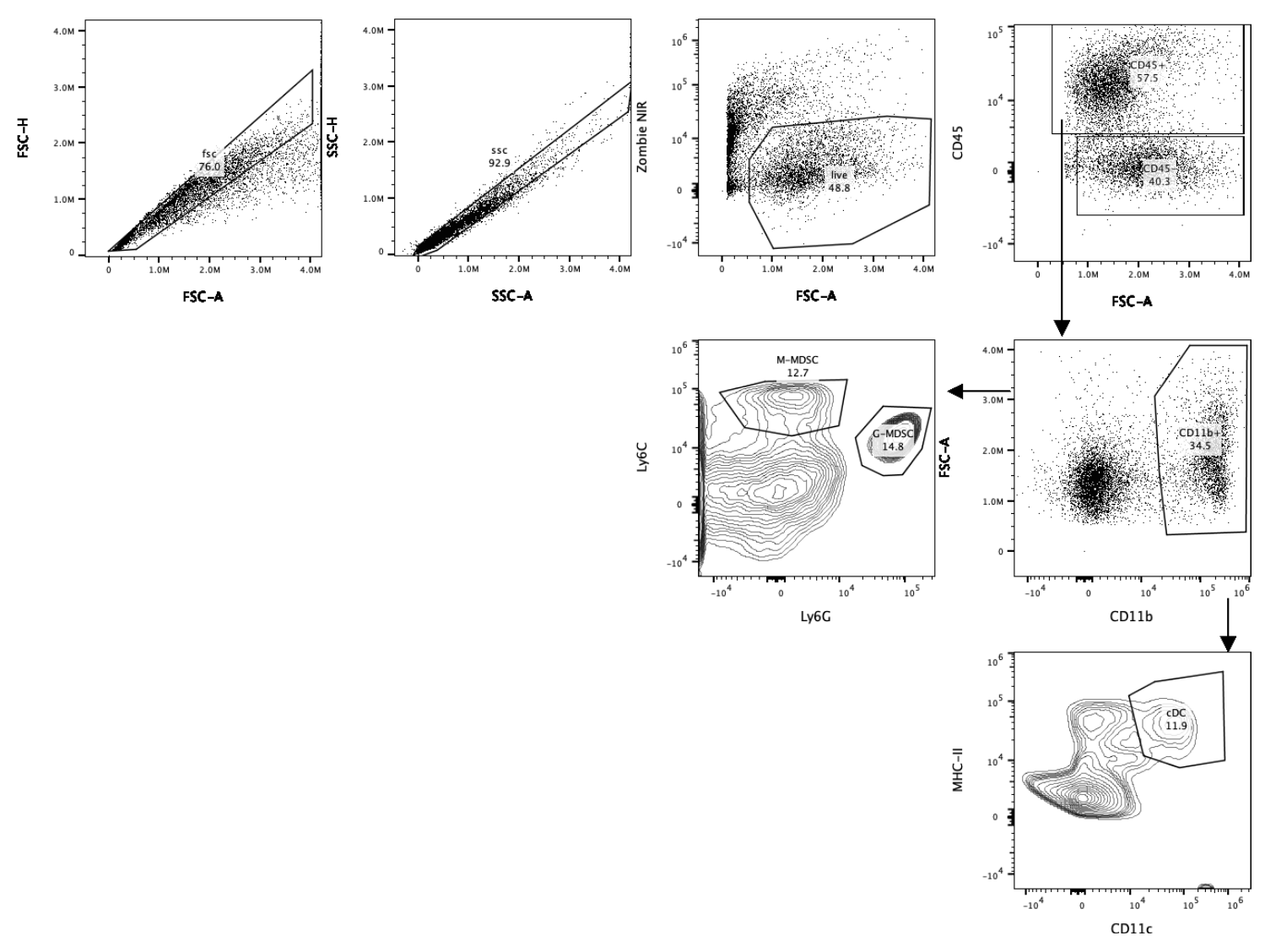

**Gating strategy of flow cytometry analysis of myeloid cell populations.** The gating example shown is on tumour cells from treated mice. After gating on singlets and cells of interest (the gate was set in the lymphocytes region using the scatter profile of the cells), leukocytes were defined by staining with anti-CD45, and then the strategy shown was used to identify different types of myeloid cells.
