## Supplementary Figures for "Tarloxotinib targets tumor hypoxia to improve therapeutic efficacy of immune checkpoint inhibition in a TLR9 dependent manner"

### Slide 1
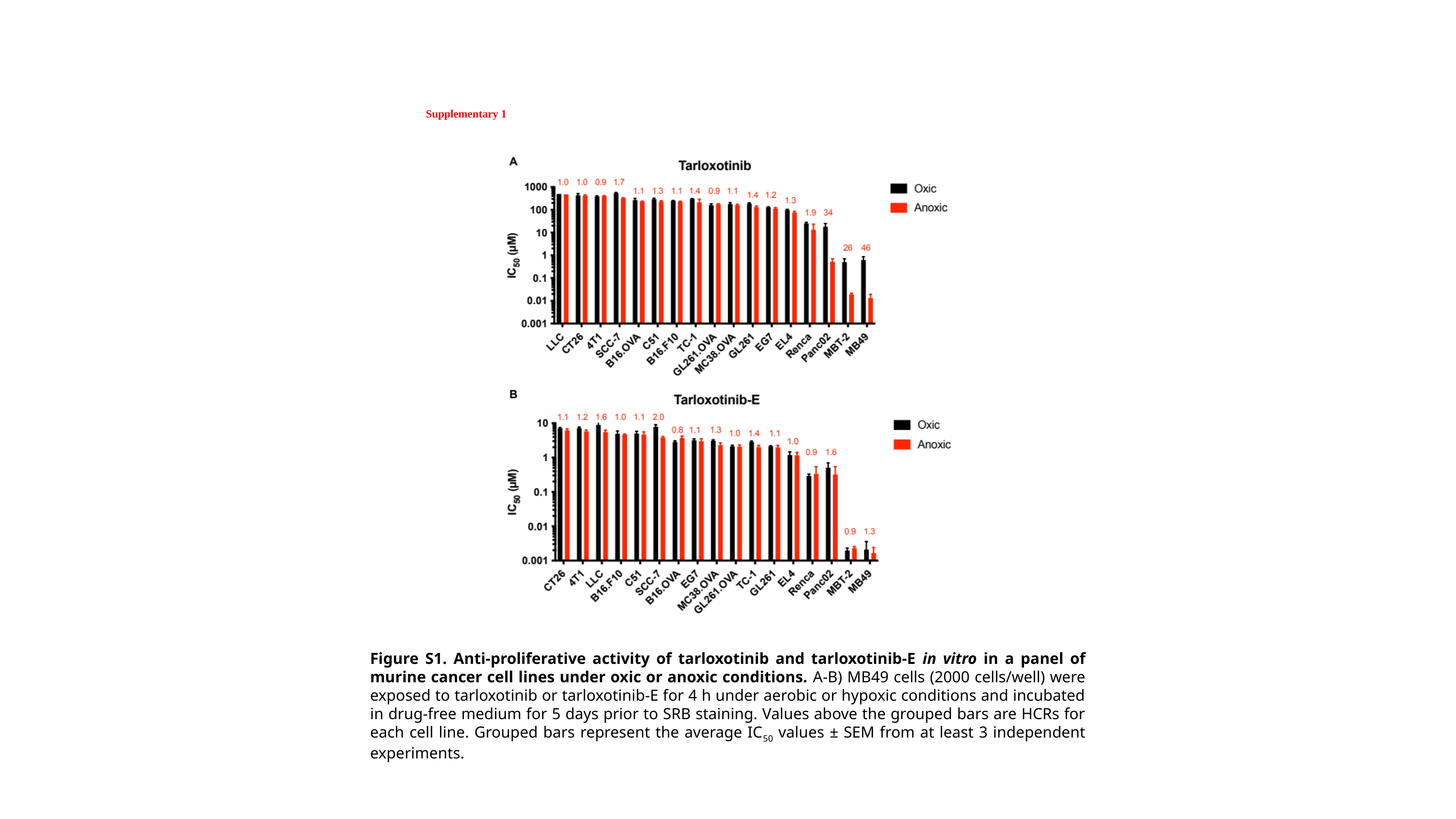

Supplementary 1
Figure S1. Anti-proliferative activity of tarloxotinib and tarloxotinib-E in vitro in a panel of murine cancer cell lines under oxic or anoxic conditions. A-B) MB49 cells (2000 cells/well) were exposed to tarloxotinib or tarloxotinib-E for 4 h under aerobic or hypoxic conditions and incubated in drug-free medium for 5 days prior to SRB staining. Values above the grouped bars are HCRs for each cell line. Grouped bars represent the average IC_50_ values ± SEM from at least 3 independent experiments.

### Slide 2
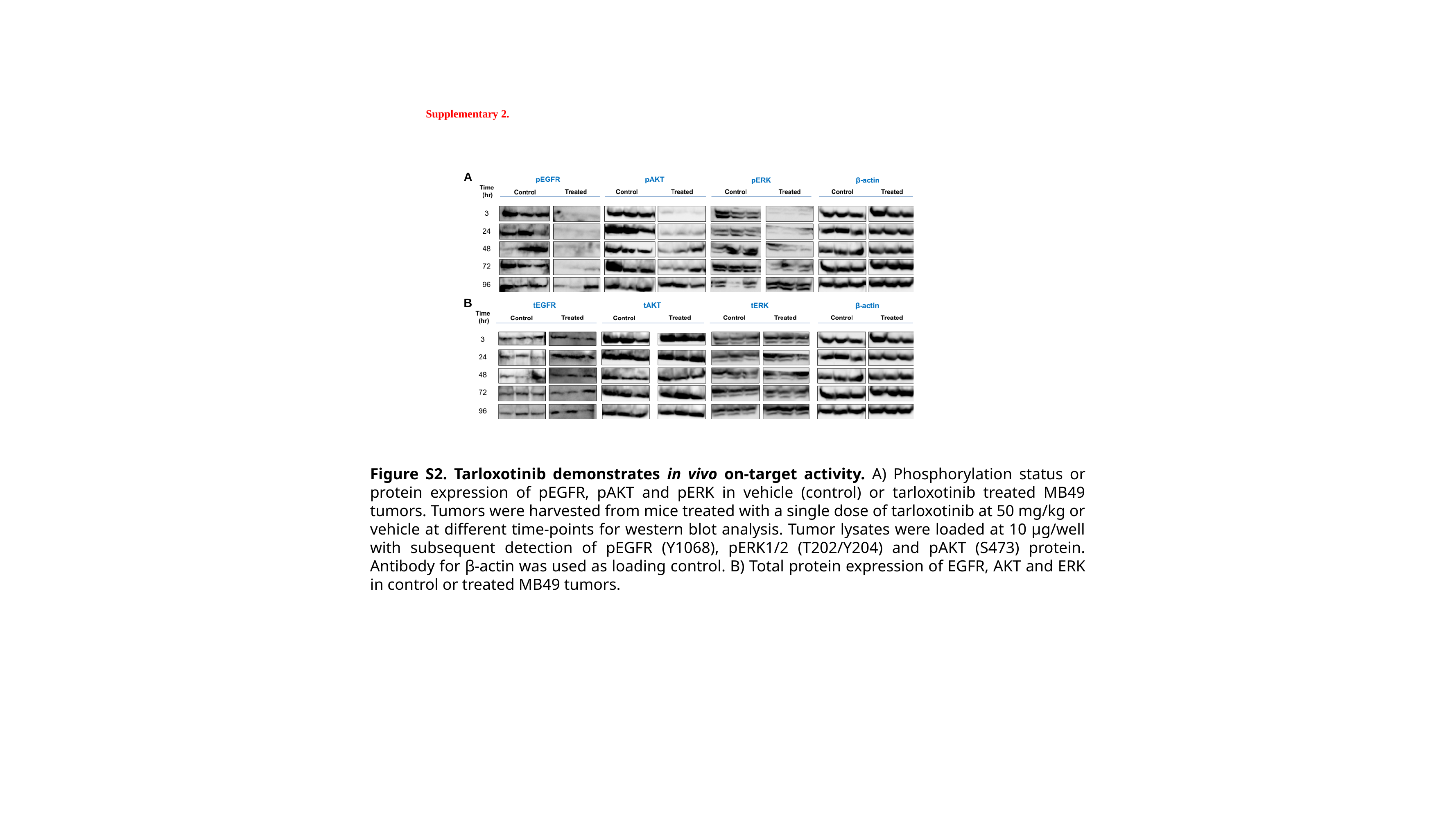

Supplementary 2.
A
B
Figure S2. Tarloxotinib demonstrates in vivo on-target activity. A) Phosphorylation status or protein expression of pEGFR, pAKT and pERK in vehicle (control) or tarloxotinib treated MB49 tumors. Tumors were harvested from mice treated with a single dose of tarloxotinib at 50 mg/kg or vehicle at different time-points for western blot analysis. Tumor lysates were loaded at 10 μg/well with subsequent detection of pEGFR (Y1068), pERK1/2 (T202/Y204) and pAKT (S473) protein. Antibody for β-actin was used as loading control. B) Total protein expression of EGFR, AKT and ERK in control or treated MB49 tumors.

### Slide 3
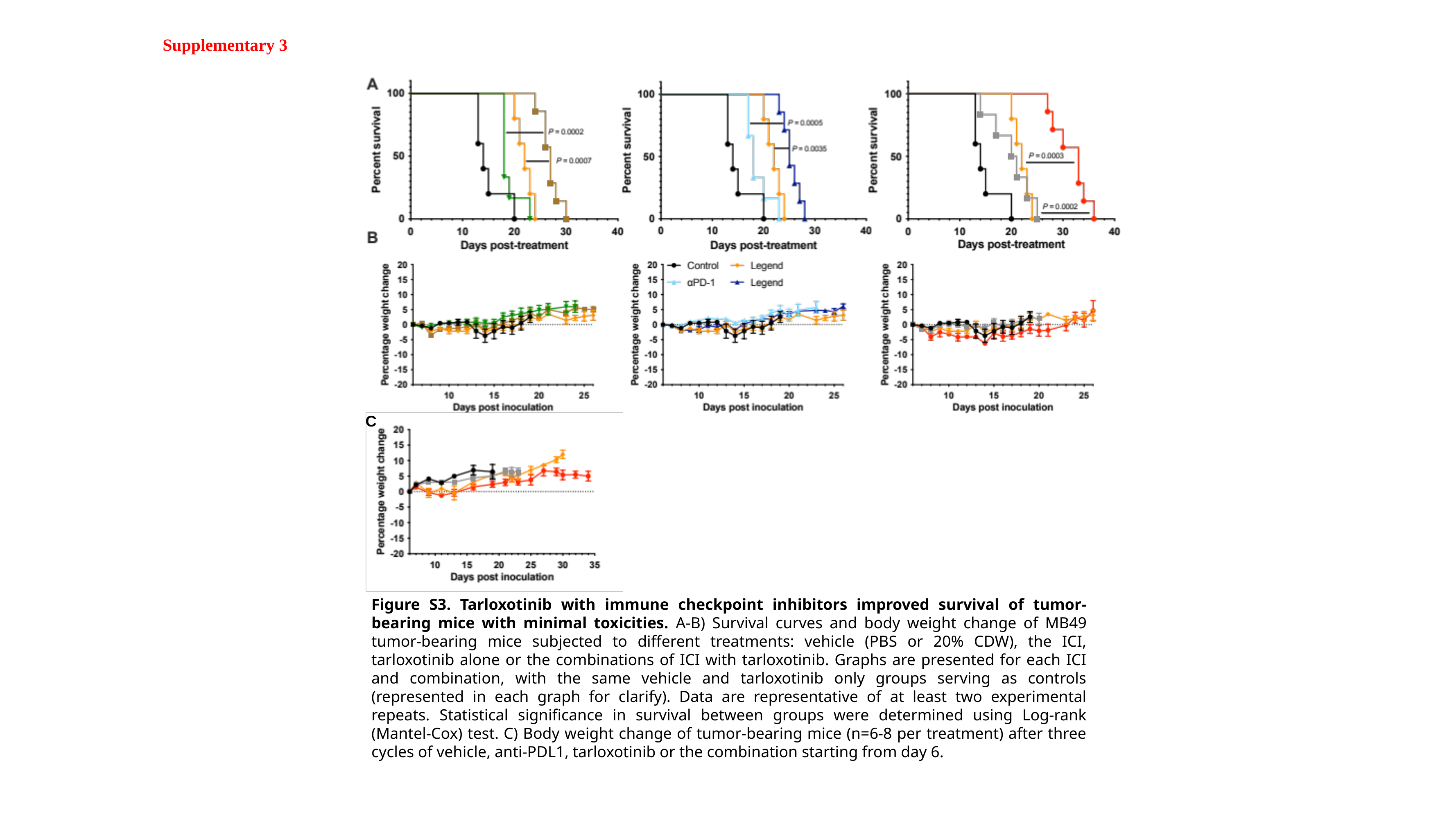

Supplementary 3
C
Figure S3. Tarloxotinib with immune checkpoint inhibitors improved survival of tumor-bearing mice with minimal toxicities. A-B) Survival curves and body weight change of MB49 tumor-bearing mice subjected to different treatments: vehicle (PBS or 20% CDW), the ICI, tarloxotinib alone or the combinations of ICI with tarloxotinib. Graphs are presented for each ICI and combination, with the same vehicle and tarloxotinib only groups serving as controls (represented in each graph for clarify). Data are representative of at least two experimental repeats. Statistical significance in survival between groups were determined using Log-rank (Mantel-Cox) test. C) Body weight change of tumor-bearing mice (n=6-8 per treatment) after three cycles of vehicle, anti-PDL1, tarloxotinib or the combination starting from day 6.

### Slide 4
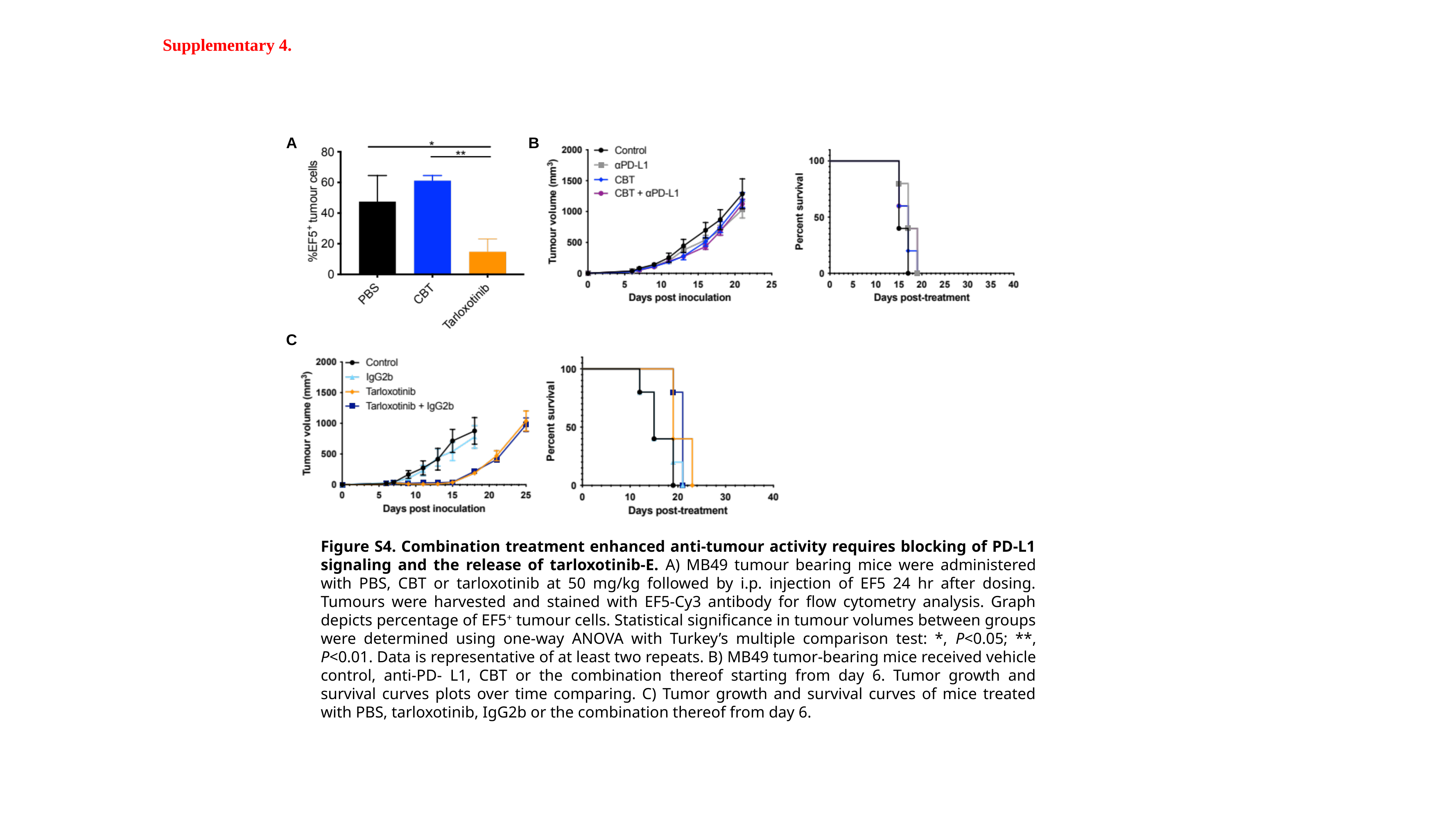

Supplementary 4.
A
B
C
Figure S4. Combination treatment enhanced anti-tumour activity requires blocking of PD-L1 signaling and the release of tarloxotinib-E. A) MB49 tumour bearing mice were administered with PBS, CBT or tarloxotinib at 50 mg/kg followed by i.p. injection of EF5 24 hr after dosing. Tumours were harvested and stained with EF5-Cy3 antibody for flow cytometry analysis. Graph depicts percentage of EF5^+^ tumour cells. Statistical significance in tumour volumes between groups were determined using one-way ANOVA with Turkey’s multiple comparison test: *, P<0.05; **, P<0.01. Data is representative of at least two repeats. B) MB49 tumor-bearing mice received vehicle control, anti-PD- L1, CBT or the combination thereof starting from day 6. Tumor growth and survival curves plots over time comparing. C) Tumor growth and survival curves of mice treated with PBS, tarloxotinib, IgG2b or the combination thereof from day 6.

### Slide 5
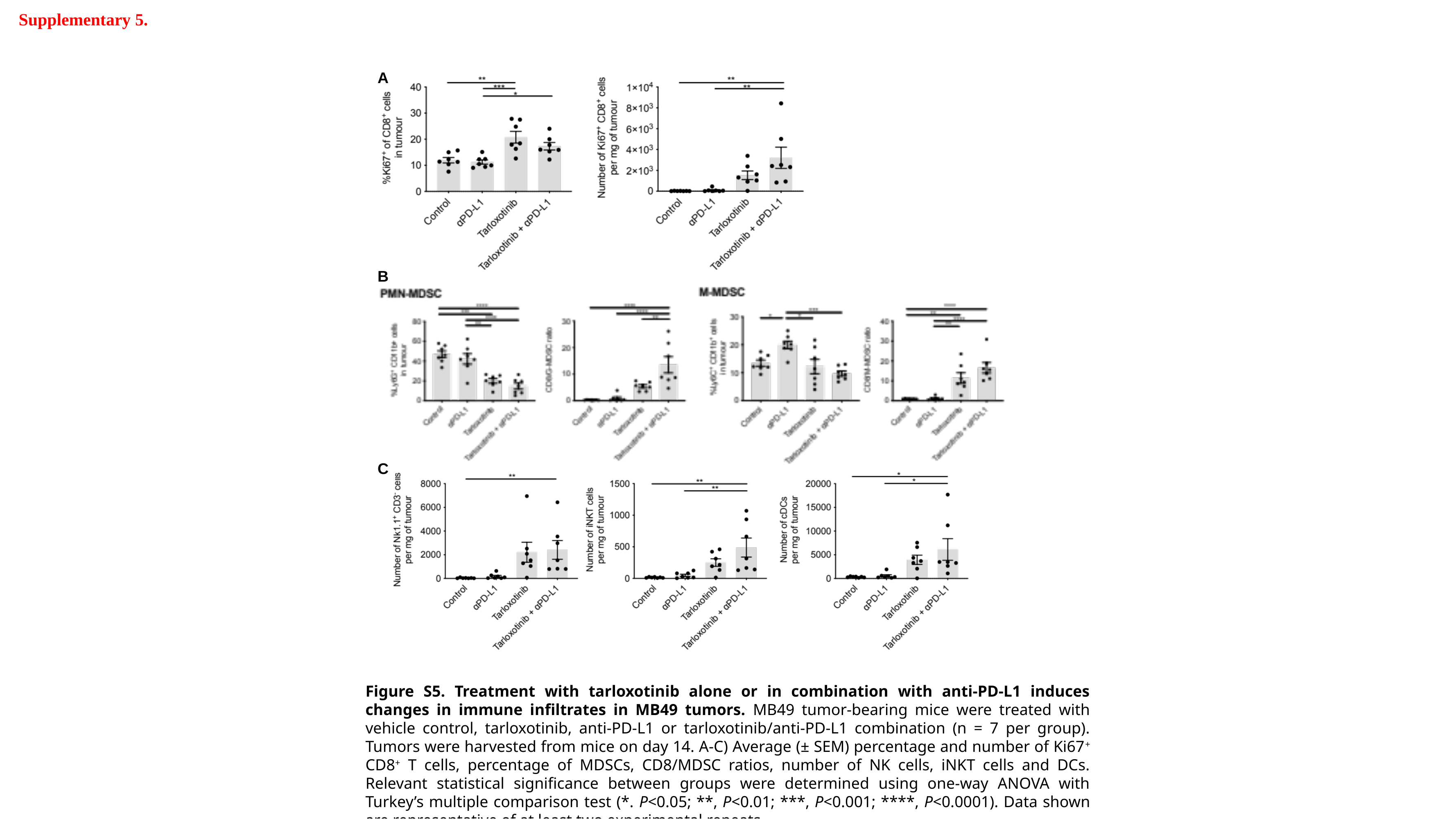

Supplementary 5.
A
B
C
Figure S5. Treatment with tarloxotinib alone or in combination with anti-PD-L1 induces changes in immune infiltrates in MB49 tumors. MB49 tumor-bearing mice were treated with vehicle control, tarloxotinib, anti-PD-L1 or tarloxotinib/anti-PD-L1 combination (n = 7 per group). Tumors were harvested from mice on day 14. A-C) Average (± SEM) percentage and number of Ki67^+^ CD8^+^ T cells, percentage of MDSCs, CD8/MDSC ratios, number of NK cells, iNKT cells and DCs. Relevant statistical significance between groups were determined using one-way ANOVA with Turkey’s multiple comparison test (*. P<0.05; **, P<0.01; ***, P<0.001; ****, P<0.0001). Data shown are representative of at least two experimental repeats.

### Slide 6
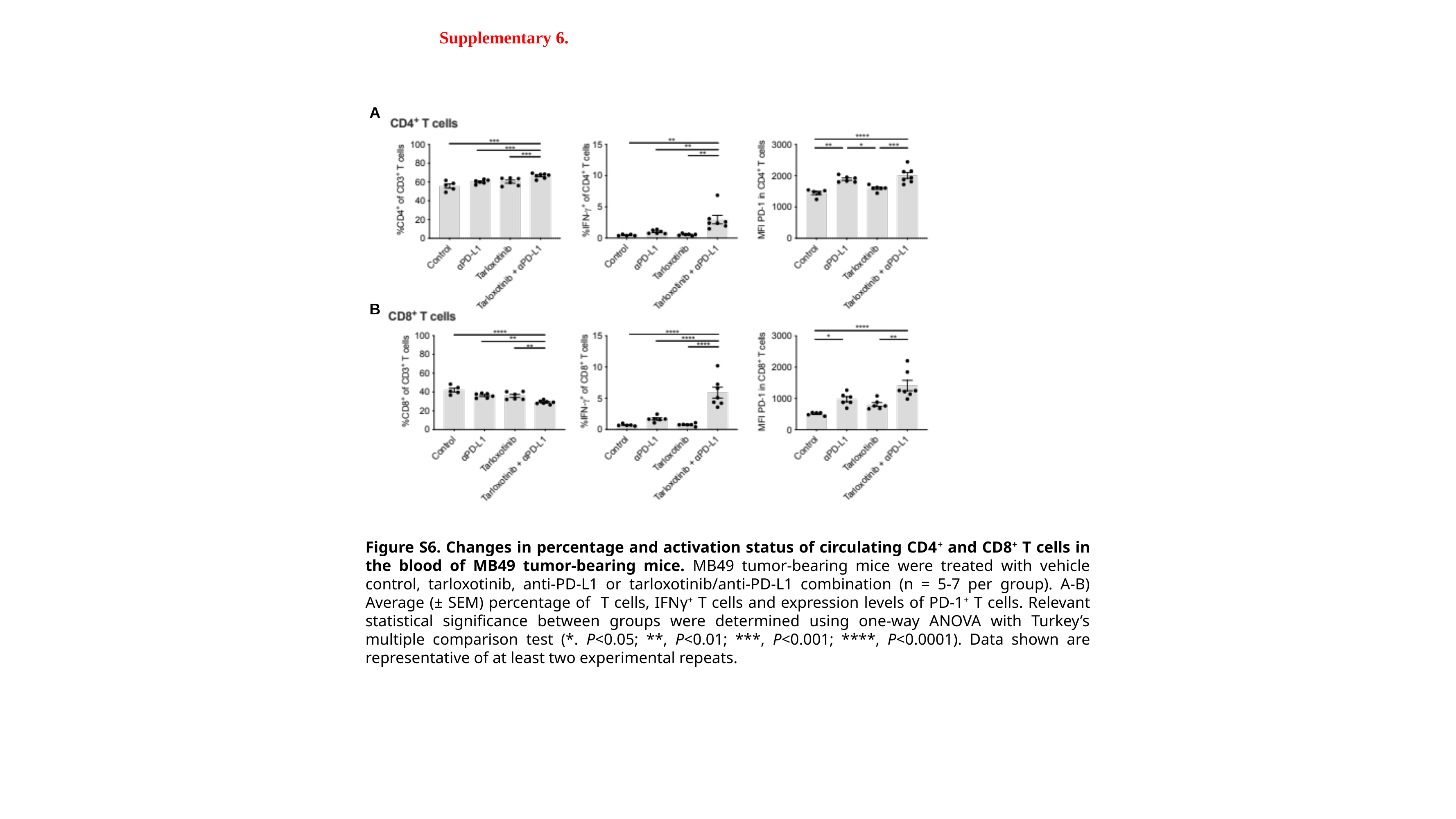

Supplementary 6.
A
B
Figure S6. Changes in percentage and activation status of circulating CD4^+^ and CD8^+^ T cells in the blood of MB49 tumor-bearing mice. MB49 tumor-bearing mice were treated with vehicle control, tarloxotinib, anti-PD-L1 or tarloxotinib/anti-PD-L1 combination (n = 5-7 per group). A-B) Average (± SEM) percentage of T cells, IFNγ^+^ T cells and expression levels of PD-1^+^ T cells. Relevant statistical significance between groups were determined using one-way ANOVA with Turkey’s multiple comparison test (*. P<0.05; **, P<0.01; ***, P<0.001; ****, P<0.0001). Data shown are representative of at least two experimental repeats.

### Slide 7
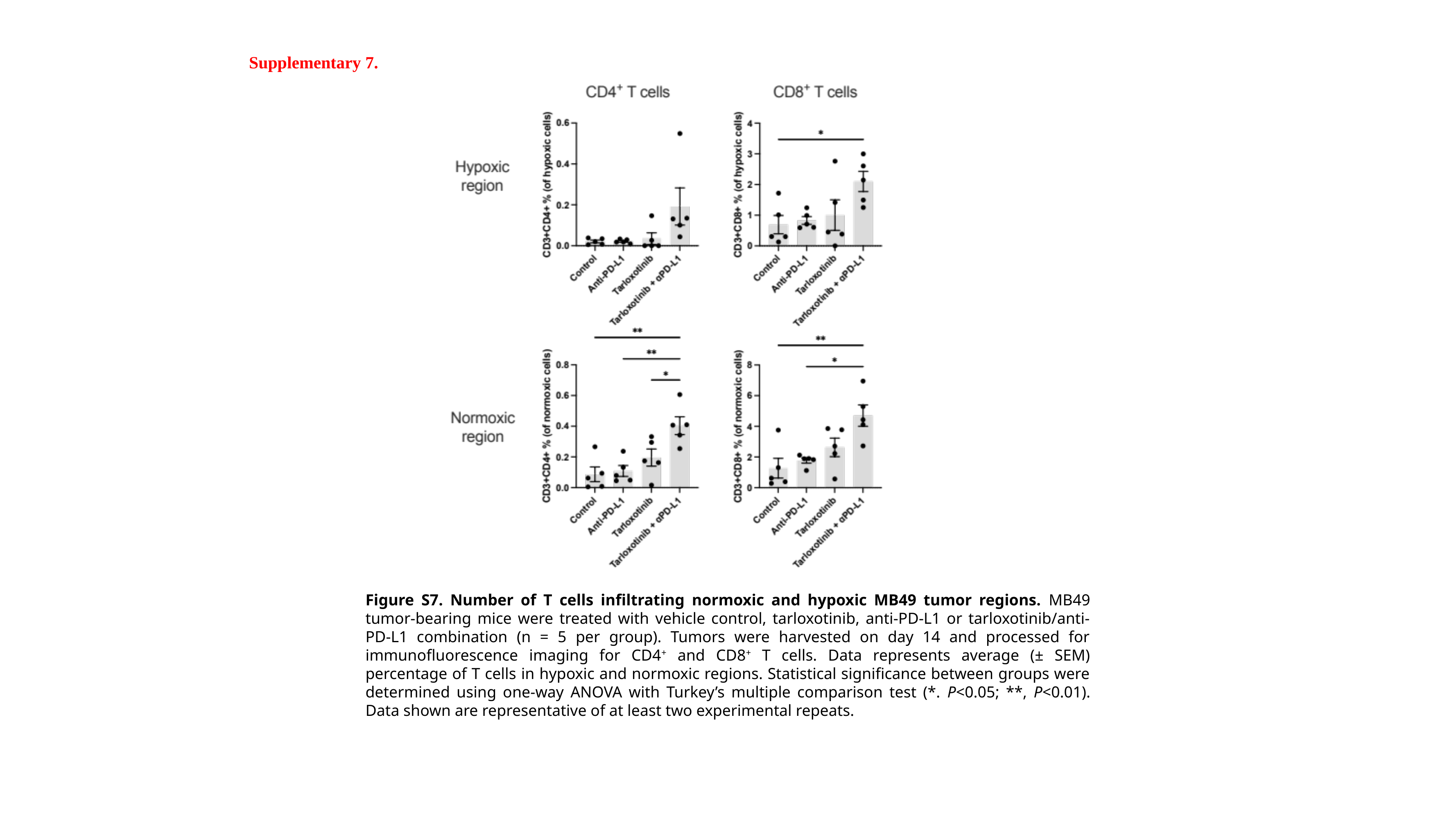

Supplementary 7.
Figure S7. Number of T cells infiltrating normoxic and hypoxic MB49 tumor regions. MB49 tumor-bearing mice were treated with vehicle control, tarloxotinib, anti-PD-L1 or tarloxotinib/anti-PD-L1 combination (n = 5 per group). Tumors were harvested on day 14 and processed for immunofluorescence imaging for CD4^+^ and CD8^+^ T cells. Data represents average (± SEM) percentage of T cells in hypoxic and normoxic regions. Statistical significance between groups were determined using one-way ANOVA with Turkey’s multiple comparison test (*. P<0.05; **, P<0.01). Data shown are representative of at least two experimental repeats.

### Slide 8
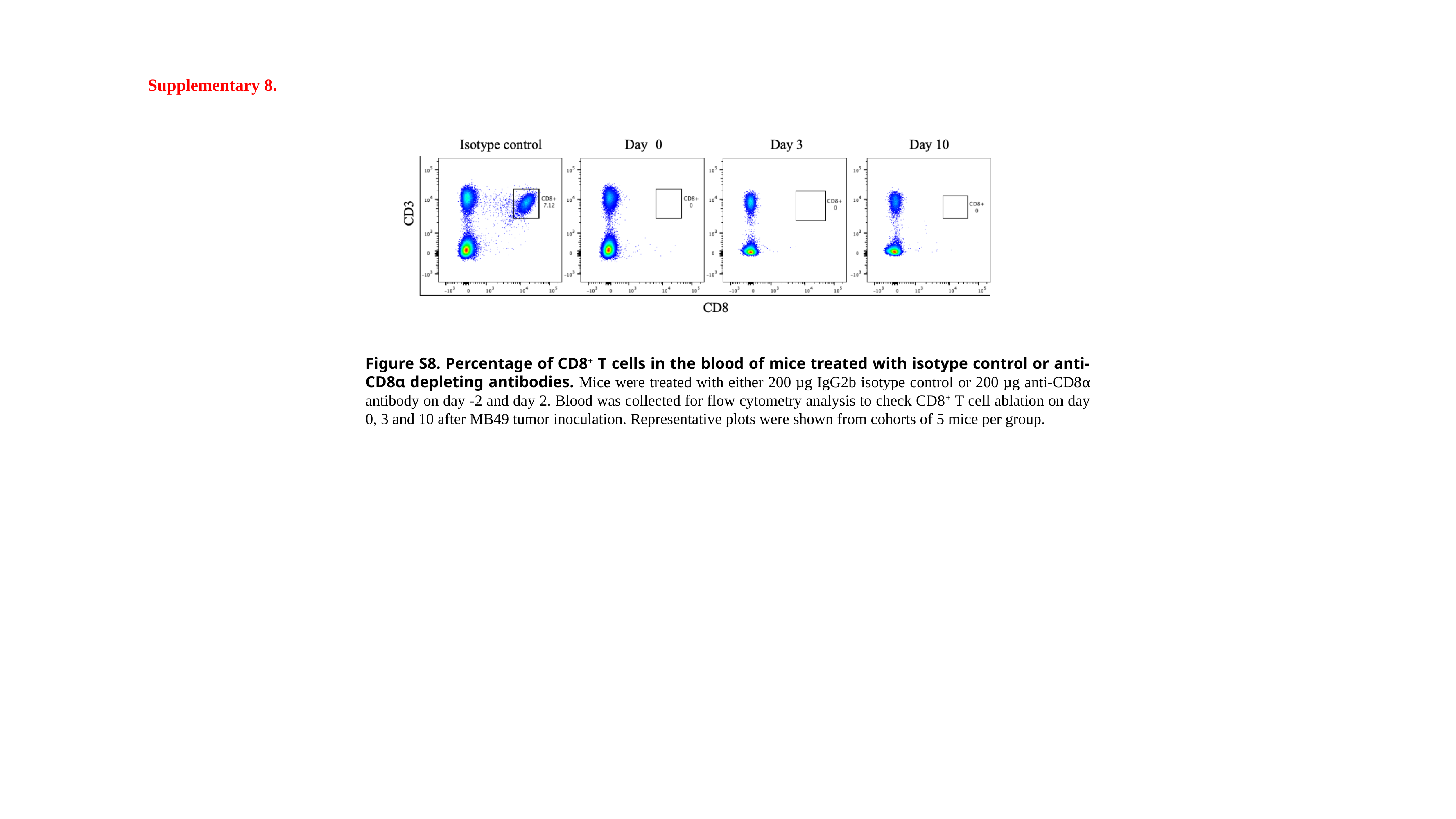

Supplementary 8.
Figure S8. Percentage of CD8^+^ T cells in the blood of mice treated with isotype control or anti-CD8α depleting antibodies. Mice were treated with either 200 µg IgG2b isotype control or 200 µg anti-CD8α antibody on day -2 and day 2. Blood was collected for flow cytometry analysis to check CD8^+^ T cell ablation on day 0, 3 and 10 after MB49 tumor inoculation. Representative plots were shown from cohorts of 5 mice per group.

### Slide 9
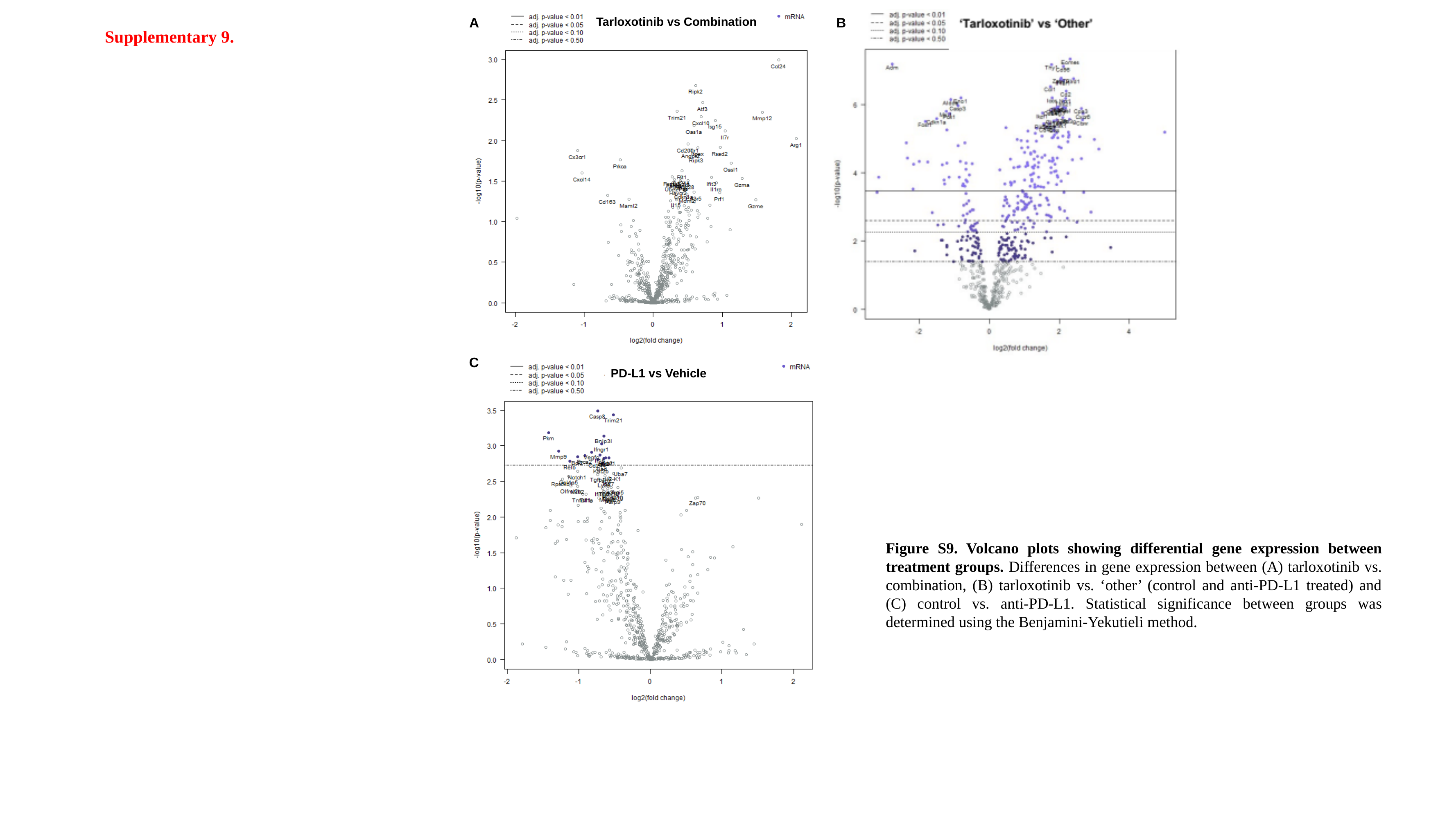

A
B
Tarloxotinib vs Combination
Supplementary 9.
C
PD-L1 vs Vehicle
Figure S9. Volcano plots showing differential gene expression between treatment groups. Differences in gene expression between (A) tarloxotinib vs. combination, (B) tarloxotinib vs. ‘other’ (control and anti-PD-L1 treated) and (C) control vs. anti-PD-L1. Statistical significance between groups was determined using the Benjamini-Yekutieli method.

### Slide 10
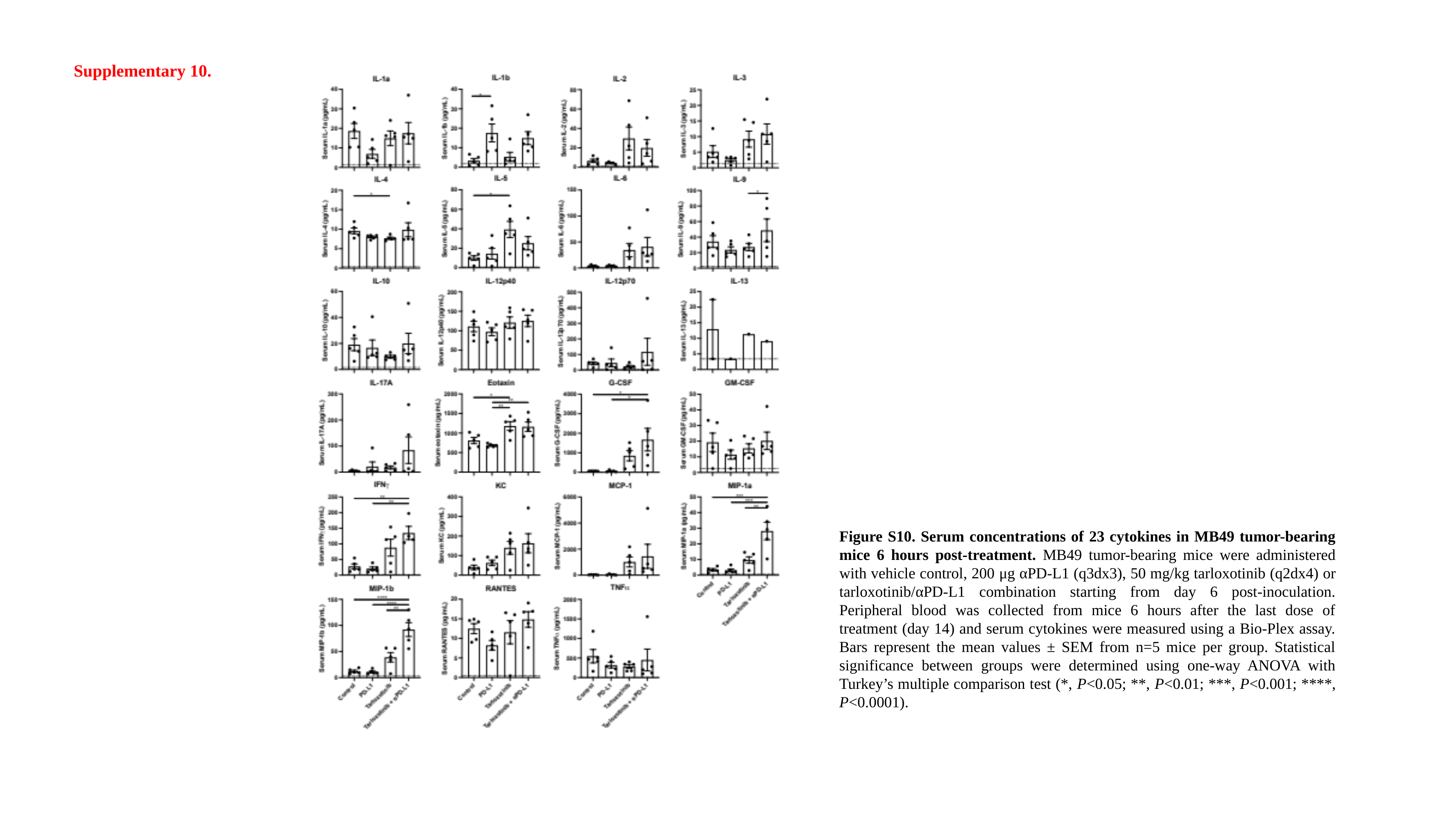

Supplementary 10.
Figure S10. Serum concentrations of 23 cytokines in MB49 tumor-bearing mice 6 hours post-treatment. MB49 tumor-bearing mice were administered with vehicle control, 200 μg αPD-L1 (q3dx3), 50 mg/kg tarloxotinib (q2dx4) or tarloxotinib/αPD-L1 combination starting from day 6 post-inoculation. Peripheral blood was collected from mice 6 hours after the last dose of treatment (day 14) and serum cytokines were measured using a Bio-Plex assay. Bars represent the mean values ± SEM from n=5 mice per group. Statistical significance between groups were determined using one-way ANOVA with Turkey’s multiple comparison test (*, P<0.05; **, P<0.01; ***, P<0.001; ****, P<0.0001).

### Slide 11
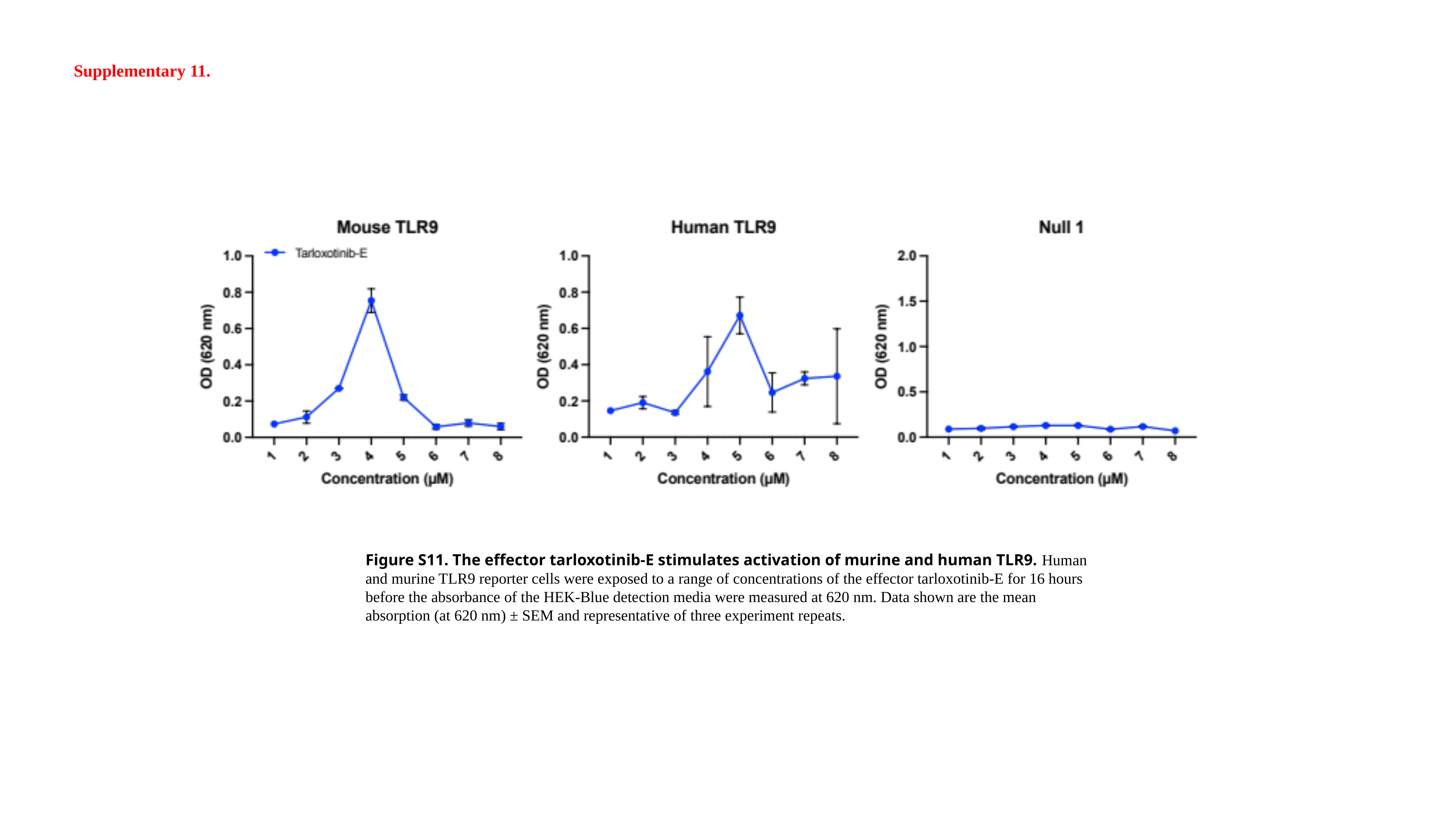

Supplementary 11.
Figure S11. The effector tarloxotinib-E stimulates activation of murine and human TLR9. Human and murine TLR9 reporter cells were exposed to a range of concentrations of the effector tarloxotinib-E for 16 hours before the absorbance of the HEK-Blue detection media were measured at 620 nm. Data shown are the mean absorption (at 620 nm) ± SEM and representative of three experiment repeats.
